# Growth Mechanics and the emergence of metabolic oscillations in growing cells

**DOI:** 10.1101/2025.06.24.661369

**Authors:** Hugo Dourado, Chilperic Armel Foko Kuate, Wolfram Liebermeister, Martin J. Lercher

## Abstract

Unicellular organisms exhibit oscillations in metabolic activity and gene expression, even when growing in static media with no external dynamic stimuli. The cause and possible function of these endogenous oscillations are still not fully understood, with different hypotheses including its origin as a byproduct of misguided regulatory processes. To investigate whether such oscillations could be functional rather than incidental, we introduce Growth Mechanics (GM) as a general mathematical framework to study the dynamic resource allocation in models of whole self-replicating cells, built exclusively on the first principles of fitness maximization, mass conservation, nonlinear reaction kinetics, and constant cell density. Inspired by the physical theory of classical mechanics, we first find the simplest mathematical description the problem in terms of generalized coordinates, and then solve the optimal dynamical resource allocation for cells growing in any given medium using the Euler-Lagrange equation, resulting in analytical “equations of optimal motion” (EOM) that apply for growing cells in general. We then solve these equations numerically for simple growing cell models and show that, in general, oscillations are necessary to maximize cell fitness by getting more saturated reactions at the time they are most needed. We also show how the EOM predict the emergence of quasi-linear dependencies of the metabolic oscillation frequency and the ribosome protein allocation with the cell growth rate at different growth media, in line with the “growth laws” observed experimentally in microbes. This work contributes to the foundations of theoretical cell biology with general quantitative principles and a bridge to theoretical tools of physics.

## I. INTRODUCTION

Oscillation is a common feature of microbial metabolism [1–4], gene expression [5, 6] and protein allocation [7, 8]. The emergence of such oscillations is currently understood in terms quantitative models for cell subsystems such as metabolism [9] and translation [5, 7], and for microbial cells as a whole by considering extra phenomenological assumptions added to the first principles that are known to shape these type of physical processes [10, 11], and as an adaptation to oscillations imposed by the growth medium, such as for photosynthetic organisms under cycles of light during day and night [12]. Currently no general quantitative theory has been able to capture, from first principles, the emergence of the oscillations also observed in microbes as a whole when growing on static media [2, 6]. The very existence of a unified theory capable of predicting general cell behavior has been questioned, since different microbes apparently employ fundamentally distinct growth strategies at different times [13]. A unifying principle common to all growing cells is however the process of evolution by natural selection, which in theoretical studies is often seen in economic terms as an optimal resource allocation problem with the objective of maximizing some measure of fitness [14–17]. The objective of this study is to address the following general theoretical questions: i) is there an absolute best strategy for growing cells to invest their resources at all times at some given dynamic growth medium, and if so, ii) does this general strategy helps explaining emergent cell behavior, such as self-oscillations. In this study we show that, under very general assumptions for microbes with no strong influence on their growth medium state, the answer is yes for both of these questions.

The main difficulty with quantitative cell models comes from the nonlinear nature of kinetic rate laws, which relate the fluxes of biochemical reactions with the concentrations of internal biomass components of the cell. This nonlinearity makes large cell models mapping the thousands of different types of reactions very challenging to treat in both analytical and numerical studies. This difficulty is partially circumvented by small nonlinear models that coarse-grain the complex reaction networks into few idealized reactions [14], and large linear models simplified by extra assumptions which make them relatively easy to be studied numerically and useful in many applications [18, 19]. The linear cell models are however still, by definition, not capable of capturing the emergence of complex behavior in a nonlinear system. These complex behavior in cells include, for example, the (apparently coordinated) continuous oscillatory resource allocation observed throughout different systems in budding yeast, including metabolism and gene expression [5, 6]. This type of behavior exemplifies how neither separate reductionist models of individual cell subsystems nor large linear models can provide the final understanding of what are the basic quantitative principles shaping cell behavior. Thus, the development of a comprehensive mathematical theory of holistic cell models as entire self-replicating systems is still a necessary step to advance our quantitative understanding of cell biology.

Self-replicating cells can be seen as complex systems involving a large network of reactions responsible for transporting chemical compounds through the cell surface, then converting these internal reactants via enzymatic reactions, until a final reaction catalyzed by ribosomes synthesizes the proteins that are distributed to catalyze all the reactions, including the protein in the ribosomes themselves. This process in growing cells may be seen as a mass flow through a hypergraph with reactions as edges and reactants as nodes [20], with the additional properties that: i) mass flow is “lost” at each node to account for the net production of each cell component (at quantities determined by mass conservation), ii) the special “ribosome reaction” must sustain the production of protein that is distributed to catalyze all reactions (at a required quantity determined by nonlinear rate laws and mass conservation), and iii) the mass concentrations of all components in the network (reactants and proteins) compose the cell density, which is finite. This finite density has in fact been observed to have an almost constant value across different bacterial species; approximately 1100 *g L*^−1^ for the buoyant density including water [21, 22] and 340 *g L*^−1^ of dry mass only [23]. The buoyant density is also almost constant during the cell cycle in many microorganisms, varying at most 0.7% in the case of the yeast *S. cerevisiae* [24].

The cell density is an important example of the still limited experimental data currently available to inform and validate quantitative cell models. Theoretical studies on the resource allocation in bacterial cells [15, 25– 30] rely on absolute transcriptomics and proteomics [25, 26, 31–33], and the much more limited absolute metabolomics [34–36] measurements for cell populations growing in different steady-state media. From these type of data, it has been long ago observed that growth rate and ribosomal protein content at different steady-state media relate in a linear way [37], a type of relationship that has been later termed “growth law” and generalized in the presence of different antibiotic concentrations [26] and for other proteome sectors [27]. Fewer experimental studies have measured how population properties change in time during shifts in growth media [38– 40], leading also to the observation of emerging nonlinear “growth laws” relating the population growth rate before and after the shift [41]. More recent technologies now allowed dynamical measurements of single bacterial cells, showing persistent fluctuations in the growth rate and ribosome allocation even during periods when the growth media is not changing [8, 42], a property which has not been reported previously at population level for bacteria possibly due to the de-synchronized growth of its individual cells [43]. Recent single-cell measurements of budding yeast have now also shown that oscillations in resource allocation can be present even when they do not show at population level [6]. While most quantitative cell models currently assume that cells growing in static growth media eventually reach an internal dynamic equilibrium state known as balanced growth, where all its extensive properties (such as biomass composition, reaction rates per volume, and growth rate) are constant in time [14–16, 30, 44], these more recent single-cell experiments indicate that oscillations are ubiquitous across unicellular organisms, and dynamical models [7, 9, 10, 45] are indeed necessary for a full understanding of the principles of cell resource allocation. Although models of cell growth at stead-state media are relevant to study lab cultures maintained at that condition, dynamical models are also a necessary step to better understand cell growth in dynamical media, which are the rule in nature.

Apart from dynamical considerations, we can also make a distinction between quantitative cell models according to their descriptive or optimization-based approach. Descriptive models focus on clarifying *how* cells do something we know they do, by taking as input all available information about the system including physical principles and phenomenological principles extrapolated from data [7, 11, 28, 40, 46, 47]. Optimization-based models focus instead on answering *what* cells would do in particular scenarios (possibly not yet tested by experiments), by searching for the optimal point maximizing some measure of fitness in the abstract space of all cell states satisfying basic constraints [14–16, 18, 19, 30, 45]. Thus, in general, optimization-based models gain some explanatory power over *what* cells would do at the cost of losing some explanatory power over *how* exactly they do it; the cell state, or its dynamical trajectory in the abstract space of possible states, is predicted and explained as the result of fitness optimization through evolution by natural selection, even if the exact mechanism realizing it remains unknown. This optimization approach is closer to how fundamental theories are currently formulated in modern physics, in favor of previously established descriptive models of specific systems. For example, early descriptive models of the solar system were replaced by Newton’s laws of motion and gravitation, which are now understood as a particular consequence of the more fundamental principle of extremal action in the later developed Lagrangian mechanics. The historical development of the many successful theories in physics into the same framework of optimizing an action functional (as defined by variational calculus) also suggests a general path for the development of fundamental quantitative theories in cell biology; but here, instead of the somewhat abstract concept of action, the very real and measurable fitness presents itself as the quantity being optimized in biology via its central concept of evolution by natural selection [48]. The formal definition of such general fitness functional depends however on an adequate optimization-based formulation for cell growth models.

Self-replicator models have been introduced by Molenaar et al. [14] as nonlinear optimization-based models of systems capable of completely replicating themselves, including the proteins necessary to catalyze all of its transport, enzymatic and ribosome reactions. It is built on the minimal assumptions of: mass conservation including dilution of all components at constant balanced growth, nonlinear rate laws, and constant protein density. At balanced growth, the growth rate is itself a direct measure of cell fitness, and its numerical optimization under these constraints was shown to be enough to explain conceptually major patterns in cell resource allocation, including overflow metabolism at faster growth [14]. Self-replicator models have also being extended to study the dynamical resource allocation in cells [45, 49], although still limited to small models due to the challenging tractability of large nonlinear optimization problems.

We have introduced previously Growth Balance Analysis (GBA) as a general mathematical framework that simplifies the description of nonlinear optimization-based models, facilitating numerical simulations and analytical derivations of general principles for cells at balanced growth [15, 16]. GBA simplifies the formulation of nonlinear cell models by i) considering a general constraint on the total cell density including all of its components, which is a conserved quantity for many microorganisms [24] and critical to capture the important trade-off between the cell investment on reactants and protein catalysts necessary to maintain reaction fluxes according to kinetic rate laws [30], ii) assuming all proteins in the cell have the same amino acid composition, and iii) a reformulation of the underlying mathematical problem on a minimal set of independent variables (one for each reaction), allowing us to use standard analytical tools (Lagrange multipliers and Karush-Kuhn-Tucker conditions) to derive general analytical conditions for optimal balanced cell growth. These fundamental quantitative principles can explain well the ribosomal protein allocation at different growth rates in the bacterium *E. coli* and the yeast *S. cerevisiae* [15]. We have also shown how this general mathematical framework helps generalizing concepts of metabolic control analysis (MCA) [50] to growth control analysis (GCA) as a theory with closed analytical expressions for how the quantitative properties of cells at balanced growth respond to small changes in their state, and how their fitness changes with small changes on cell density, growth medium concentrations and kinetic parameters [15, 16]. In this study we focus on generalizing these previous analytical results on optimal balanced growth states [15, 16] to optimal dynamical states of cells growing in any given growth medium. We then use these results to show how self-oscillations always emerge in optimally growing cells entirely from first principles, generalizing previous theoretical studies on how metabolic oscillations emerge under extra simplifying assumptions [9, 10], or forced by internal enzymatic oscillations [51]. We introduce now some key mathematical concepts necessary to define the optimization problem we want to solve.

### A. Growth rate and fitness

A central concept in this study is of the *instantaneous growth rate µ*(*t*) (also known as specific growth rate, and hereafter referred simply as growth rate for short), here defined as the instantaneous proportional change in the mass *m*(*t*) of a cell at time *t*

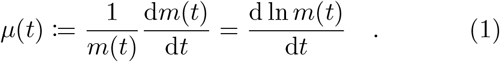

At constant cell density *ρ* (mass per volume in *g L*^−1^ units), the definition (1) above is also equivalent to the instantaneous proportional change in cell volume [9, 14]. A growing cell eventually replicates by dividing into two similar copies of itself, a cumulative process resulting in a population of similar cells. For simplicity, we continue thinking of this growing population of similar cells as one “big” cell that is growing in mass with growth rate also defined by equation (1). The fitness of this cell (population) during a time *T* is quantified by its *average growth rate* [9, 45]

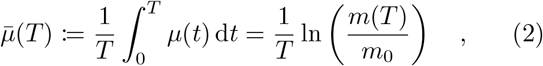

where *m*_0_ is the initial mass. The average growth rate (2) is a direct measure of fitness since the populations with higher 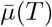 have the larger proportional growth in size (here quantified by mass) over competitors at the same period *T*, thus it conveys enough quantitative information to determine which populations are favored by natural selection. Another related concept is the long term growth rate Λ, defined as the limit of the average growth rate at infinite time, if it exists [9, 10, 48]

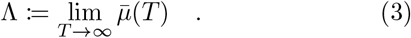

At balanced growth, the growth rate (1) is constant in time, so from equation (2) we have that *µ* becomes itself an equivalent measure of fitness [14]. Previous theoretical studies on cells out of balanced growth have found, under extra phenomenological assumptions, solutions for the optimal cell resource allocation maximizing the instantaneous growth rate (1) or the long term fitness (3) as proxies for fitness, but not the actual fitness at finite time periods *T* as quantified by the average growth rate (2). Here we solve analytically the optimization of average growth rate at arbitrary *T* for growing cells in general, entirely from first principles as formally defined next.

## II. RESULTS

In the following we first formalize the general optimization problem underlying the growth of cells in any given medium, and then present a reformulation for this problem in terms of generalized coordinates and their corresponding generalized velocities. This reformulation allows us to directly use the Euler-Lagrange equations to derive the “equations of optimal motion” (EOM) determining the dynamic resource allocation of any optimally growing cell. We then present examples of numerical solutions for EOM in static and dynamical media, showing that oscillations emerge as a way to maximize cell fitness.

### A. Cell growth models (CGMs)

To solve the mathematical problem we are interested, we first formalize the concept of cell growth models (CGMs), which captures all relevant properties of growing cells we are interested. As for self-replicator models [14], we consider models that can be seen as cellular reaction networks equipped with two extra properties: i) complete self-replication, in the sense that each reactant in the network (i.e. each node) has an associated net production flux “taking” mass flux at that node to balance the dilution by growth and concentration change of the corresponding reactant, and ii) having one “ribosome” reaction producing the proteins – all assumed to have the same composition, for simplicity [14, 16, 26, 45, 47] – that catalyze each of the reactions (hyperedges in the network, if it is seen as a hypergraph). Because all proteins are assumed to have the same composition, we simplify the following mathematical formalism by accounting for their net production flux only once in a single node “p” representing the proteome as a whole. This simplification on the proteins composition avoids doubling the number of necessary variables to characterize cell states [44], and it is justified by the minimal variation in amino acid composition of the *E. coli* proteome across various conditions [16, 31]. The resulting cellular reaction networks enriched with those two properties are now a type of cell models with the important feature of not requiring phenomenological assumptions about the cell biomass composition, which are otherwise needed as model inputs in other modeling frameworks [18, 19]; here, this composition emerges instead as an output of the growth optimization process, enabling the model to capture global trade-offs in optimal cellular resource allocation from first principles only [14]. Differently from self-replicator models, we substitute the constraint on the total protein concentration by a constraint on the total cell density, including proteins and all other non-protein components. The following detailed definition has been introduced by us before in the context of growth balance analysis (GBA) [15], but here we update this terminology to emphasize how this same mathematical object (CGM) is relevant to study cell growth in general, not only at balanced growth.

We define a cell growth model as the mathematical object completely determined by the three inputs (**M**, *ρ*, ***τ***) with the following properties:

- The mass fraction matrix **M** with real entries quantifying in each column *α* the mass fraction of each internal reactant (in a corresponding row) being consumed by it (if negative), produced by it (if positive), or neither (zero entry). We reserve by default the first columns of **M** to be (at least one) transport reaction(s) which exchange mass with the growth medium, followed by columns of enzymatic reaction(s) which convert only internal reactants, and the last column r to the ribosome reaction converting internal reactants into protein, by default in the last row p; so the last row contains only zeros and the entry 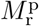, which corresponds to the mass fraction of the ribosome reaction flux that is going into proteins, as shown below

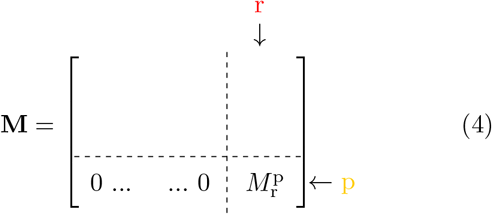

Because the entries in each column of **M** are the mass fractions of reactants going into each reaction (negative entries), going out of reactions (positive entries) or neither (zero entries), the columns corresponding to enzymatic and ribosome reactions must sum to zero due to the mass conservation of internal reactants. Transport reactions exchange mass with the growth medium, so by definition their corresponding column sums must be different than zero. This column sum property will be later relevant in our derivations.
- The cell density *ρ* in *gL*^−1^, which we assume is constant in time 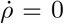 and distributed across the cell biomass at mass concentrations *ρ* **b**, where **b** is an adimensional vector of biomass fractions that sum to one.
- The vector ***τ*** = ***τ*** (**a**, *ρ* **b**) of turnover times (in *h*) encoding the rate law (typically the Michaelis-Menten rate law) for each reaction as a function of the vector **a** of reactant concentrations in the growth medium, and the concentrations of internal reactants *ρ* **b**. More specifically, i) only the turnover times of transport reactions depend on the external reactant concentrations **a**, and ii) each turnover time *τ*_*α*_ depends the concentrations of few reactants only: the substrates consumed by the reaction *α*, and possibly also its products and other few reactants that may act as inhibitors or activators. Note that by the nature of these physical processes, kinetic rate laws depend on the concentration of reactants (abundance per cell volume, here in *gL*^−1^) not simply abundances (e.g. in *g*/cell or *mol*/cell). The turnover time *τ*_*α*_ of each reaction *α* quantifies the mass of substrates consumed per mass of protein catalyst (transporter, enzyme or ribosome) per hour, so it relates the flux *v*_*α*_ (in *gL*^−1^*h*^−1^) with the necessary fraction *ϕ*_*α*_ of the total protein concentration *b*_p_ *ρ* allocated to catalyze the reaction

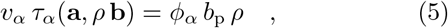

where protein allocations ***ϕ*** sum to one by definition, and *b*_p_ is the mass fraction of protein in the biomass. In general, reactions get continuously more saturated at higher concentrations of their substrates (the reactants consumed by them), meaning turnover times are smooth (non-constant) functions that converge to a nonzero value as substrate concentrations become very high. This effect can only be captured by nonlinear rate laws, the most common being the Michaelis-Menten kinetics (see SI Appendix kinetic rate laws). The saturation of reactions impose a fundamental trade-off in cell economics: for reactions going forward (i.e. positive flux *v*_*α*_ and turnover time *τ*_*α*_), higher substrate concentrations mean a lower turnover time *τ*_*α*_, so according to equation (5) the cell can maintain the same flux *v*_*α*_ with less protein *ϕ*_*α*_ allocated to this reaction, saving its valuable protein resource *b*_p_ *ρ*.

We have shown before that the trade-offs in cell economics originated from saturable (i.e. nonlinear) kinetic rate laws and limited cell biomass are critical to understand the optimal balance of enzyme and substrate allocations, which is closely implemented in real bacterial cells [30]. These trade-offs will also be central in the understanding of our results on optimal dynamical cell states, as defined next.

### B. Growth states and growth trajectories

Not all possible combinations of cell state properties such as growth rate *µ*, reaction fluxes **v** and biomass fractions **b** represent valid states for growing cells. For a given CGM (**M**, *ρ*, ***τ***) and medium defined by a function **a**(*t*) determining its composition in time, these variables must satisfy the important physical constraints of mass conservation (including dilution by growth and concentration changes at constant cell density *ρ*) for a given initial biomass composition **b**(0) = **b**_0_

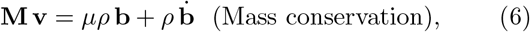

kinetic rate laws for all reactions, which we can encode in one single equation by summing equation (5) for all reactions *α* considering that the proteome fractions sum to one ∑ ***ϕ*** = 1

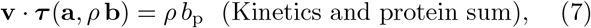

and that biomass fractions sum to one

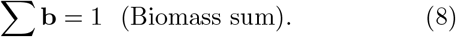

For a given CGM and medium **a**(*t*), a growth state at a point in time *t* is as a combination of variables 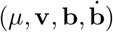 satisfying the equality constraints (6,7,8) and the inequality constraints of non-negative: biomass fractions **b**, proteome fractions ***ϕ***, and ribosome fractions ***χ*** allocated to the production of each protein

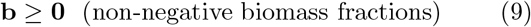

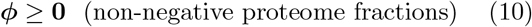

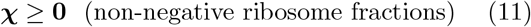

where the last two inequalities (10,11) may be equivalently stated in terms of the variables 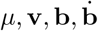, (see methods). Note we do not impose inequality constraints on fluxes **v**, as it is usual in linear cell models to partially account for the thermodynamics of reactions [18, 19]; here the thermodynamics involved in each reaction is fully captured by the nonlinear functions ***τ*** and the inequality on proteome fractions ***ϕ*** (10), which enforce that for each reaction *α* its flux *v*_*α*_ and turnover time *τ*_*α*_ must have the same sign (see equation 5, where *ρ, b*_p_ are non-negative by definition and by inequality (9), respectively). In this study, we focus on the equality constraints (6,7,8) since these are always active and relevant to study optimal growth states, while the inequality constraints (9,10,11) are in many cases inactive, meaning they do not constrain these optimal states. We thus develop in the following study an analytical theory that accounts only for the equality constraints, and check a posteriori in numerical solutions when the inequality constraints are also satisfied. On Fig.(1) we present the example of a simple CGM “L3”, and its corresponding mathematical expressions determining possible growth states.

**FIG. 1.**
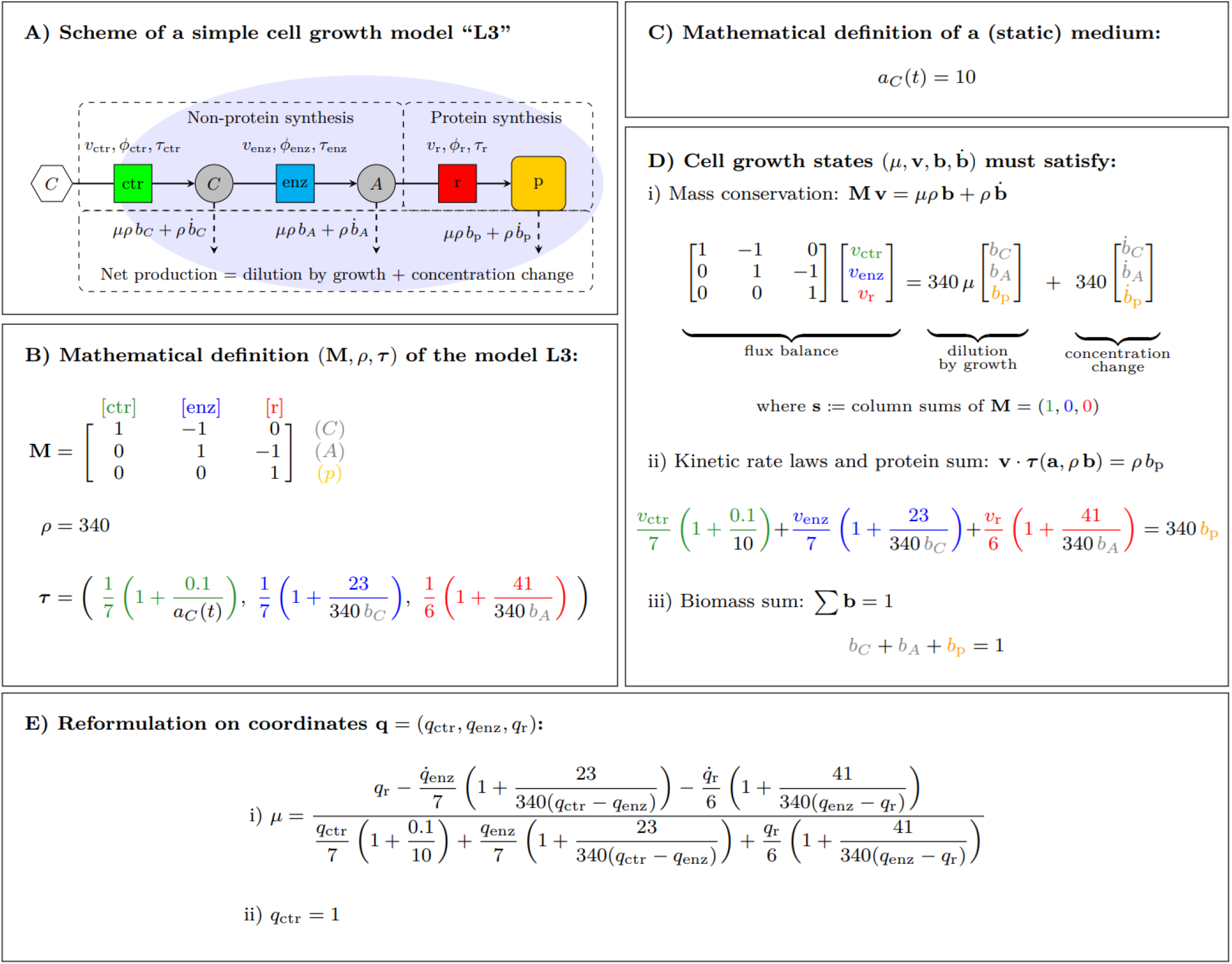
A simple cell growth model “L3” with three reactions (solid arrows, labeled by squares) in a linear pathway, and the corresponding mathematical expressions determining possible growth states. **A)** Model scheme representing the main processes defining a cell growth model: i) a “non-protein synthesis” subnetwork exchanging reactants with the growth medium (in this simple case, only one reaction “ctr” transporting the reactant “C” representing carbon) and synthesizing the necessary reactants (in this case, via one enzymatic reaction “enz” producing “A”, representing amino acids) required for ii) the “protein synthesis” via a ribosome reaction “r”, at a rate necessary to satisfy at each point in time iii) the net production of protein due to the dilution by growth and the concentration change (dashed arrows) of proteins in the model (here “ctr”, “enz” and “r” have the same composition and are thus accounted only together by their total sum “p”), which we also account for all internal reactants (gray circles) in the model, depending on the growth rate *µ*, the given cell (constant) density *ρ* and biomass fractions *b* at each point in time. In other modeling frameworks, this net production is simplified by assuming that only proteins need to be produced for the cell to grow [14, 26, 45], or a reduced set of proteins and important reactants needs to be produced via a phenomenological “biomass reaction” [19, 52–54]. **B)** A cell growth model is mathematically defined by only three inputs: i) the mass fraction matrix **M** encoding the structure of the reaction network and the mass conservation within reactions, ii) the constant cell density *ρ* in *g L*^−1^ encoding the limit on internal concentrations and iii) the function ***τ*** = ***τ*** (**a**, *ρ* **b**) encoding the kinetic properties of the reactions, in this example following the Michaelis-Menten kinetics (see Appendix). **C)** The growth medium is uniquely determined by a given function **a**(*t*), encoding how the medium concentrations change in time (in this simple example, a single concentration *a*_*C*_ = 10 defines a static medium). **D)** The possible growth states 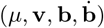 for the model defined in (B) and medium defined in (C) must satisfy the constraints of i) mass conservation accounting for dilution by growth and concentration changes for all cell components (with the mass balance within reactions encoded in the vector **s** defined as the column sums of **M**, internal reactions conserve mass and must have zero entries in the vector **s**), ii) kinetic rate laws combined with the condition that the ribosome reaction produces the sum of all proteins, and iii) the biomass fractions sum to one. **E)** In the following results, we show how all the implicit constraints on the growth rate *µ* (for this CGM, the 5 equations on 9 variables shown in D) can be reformulated on an explicit equation for the growth rate on “generalized coordinates” **q** (in this case, three) and a single continuity constraint on the transport reactions (in this case, *q*_*trs*_ = 1).

We say a growth state with 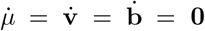 is a balanced growth state (BGS), and the balanced growth state (*µ*, **v, b**) with maximal growth rate *µ*^*\**^ is the optimal growth state (OGS) (*µ*^*\**^, **v**^*\**^, **b**^*\**^) [15, 16]. The previous definitions of growth state, BGS and OGS refer to cell states at a given point *t* in time; we now define the more relevant concept for this study on the dynamics of cell growth: for some given CGM, medium **a**(*t*) and time *T*, a growth trajectory 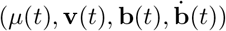 associates each point in time *t ∈* [0, *T*] to a valid growth state 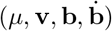. The main focus of this study is then on the optimal growth trajectory (OGT), defined here as the growth trajectory 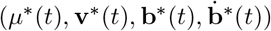 with maximal fitness, quantified by the average growth rate (2). We now redefine both growth states and trajectories in terms of much fewer variables, which will allow us to solve the optimal dynamical resource allocation problem by using the Euler-Lagrange equation.

### C. Reformulation on generalized coordinates

Our key contribution in this study is the introduction of the adimensional “generalized coordinates” vector **q** = **q**(*t*) satisfying the following equation for the biomass fractions **b** = **b**(*t*) at all points *t* in time

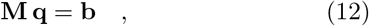

which means their time derivatives also relate as

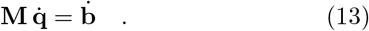

We refer to 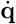 as the “generalized velocities”, in analogy to classical mechanics. The vector **q** always exists for any given initial biomass **b**_0_ if **M** is full row rank (i.e. has linearly independent rows). More generally, **q** always exists if **b**_0_ satisfies conserved moieties (i.e. **b**_0_ *∈* Im(**M**)).

Now substituting both 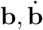 on the mass conservation equation (6) with equations (12,13), respectively

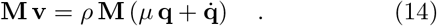

If the matrix **M** is full column rank (i.e. has linearly independent columns), **v** = **v**(*t*) is uniquely determined as

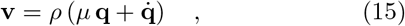

and satisfies mass conservation (6) by construction. For simplicity in our study focused on general properties of growing cells, we continue our derivation assuming that **M** is full column rank, which is effectively the case in active reaction networks at optimal balanced growth [15, 44], and more generally can always be achieved by enough coarse-graining when creating a CGM; one can always “lump” linearly dependent columns of **M** as single idealized reactions, so our following results still capture the general properties of growing cells modeled in this way. A possible extension of this present work for general matrices **M** may consider a similar approach as we used before for optimal balanced growth via KarushKuhn-Tucker (KKT) conditions [16].

We now use equations (12, 15) to substitute **b** and **v** in equation (7), respectively

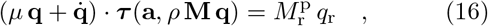

with a factor *ρ* canceling on both sides, and the RHS reminiscent from the substitution 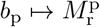 (the ribosome reaction is the only reaction producing proteins). The factor 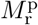 corresponds to the mass fraction of the ribosome reaction flux that is going into proteins, an adimensional factor that will carry on in later expressions as a sort of “fundamental constant” in our systems. Now we solve equation (16) for *µ*

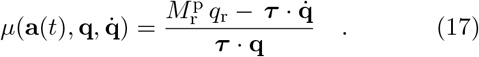

The only remaining constraint to be accounted for is the density constraint (8), which we can rewrite in terms of **q** via equation (12)

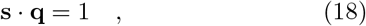

where **s** is the vector defined as the column sums of **M**, which has non-zero entries only for transport reactions since these are the ones which do not conserve internal mass (see SI Appendix, column sum vector **s**), thus equation (18) can be seen as a type of continuity equation on the transporters. Note that in the simplest case of models with only one transport reaction, equation (18) already determines a constant value for the first entry in the vector **q** corresponding to the transport reaction (as in the example in Fig.(1)), and by consequence the first entry in the generalized velocities 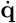 would be zero.

Our system of interest is now completely determined by **q** (including the functions ***τ***) and its time derivative 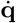. We could still go further and formulate the problem on truly independent generalized coordinates by incorporating the continuity constraint (18) into a single equation for the constrained growth rate, but this is not strictly necessary for the main purposes of this study and it is left for a future development of this work.

We define now a Lagrangian function ℒ encoding the constrained growth rate as a function of 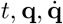

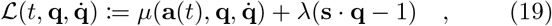

where *λ* is a Lagrange multiplier. That was the last step necessary to now being able to formally define a general functional for the fitness of growing cells.

### D. The principle of extremal fitness

We now define the *average growth functional*, 𝒜 quantifying cell fitness over a time *T* for a given CGM (**M**, *ρ*, ***τ***) and growth medium **a**(*t*) as

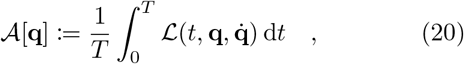

where the function 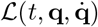 is given by equation (19). Equation (20) can be seen as an update on the original equation for the average growth rate (2), but now including all the information about constraints on growth and with an explicit dependence on 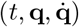. From this point on, we arrived at the same kind of mathematical framework employed in fundamental theories in physics: once all the information about the system of interest is encoded into a single function (the Lagrangian), we can find the differential equations governing the dynamical behavior of our system by solving the corresponding Euler-Lagrange equations, which guarantee that the associated functional defined as the integral of this Lagrangian in time (the action in physics, here the average growth rate *A*) is extremal (i.e. *δA* = 0).

### E. The equations of optimal motion (EOM)

Calculating now the Euler-Lagrange equations, we get for each reaction *α*

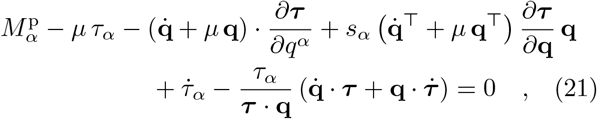

where *µ* is given by equation (17) and the derivatives of ***τ*** are calculated according to the chain rule, with *∂****τ*** */∂q*^*α*^ seen here as a vector and *∂****τ*** */∂***q** as a matrix (see SI Appendix). The last equation (21) together with the continuity constraint (18) determine implicitly the velocities 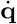 in terms of some functions on time *t* and coordinates **q**, via a system of nonlinear first order ordinary differential-algebraic equations (DAE) for the optimal growth trajectory (OGT) of some given: i) CGM (**M**, *ρ*, ***τ***), ii) growth medium **a**(*t*) and iii) initial conditions 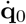. We thus name (18,21) the equations of optimal motion (EOM), in analogy to the equations of motion in physics, but emphasizing that these apply only to the hypothesized *optimal* growth trajectories **q**^*\**^(*t*). The equation (21) for steady-state initial conditions and medium 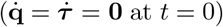 reduces to the system of algebraic equations we found before [16] for optimal growth states **q**^*\**^ in static media, indicating the regime where our previous results on optimal balanced growth are still relevant.

In this work we focused on finding the best possible resource allocation strategy for cells growing on a predefined dynamic medium, and showed that this hypothetical strategy must satisfy equations (21,18), if it exists; one can however always construct simple counterexamples of “faulty” GCMs that cannot growth in the first place, so there is always the possibility of no solutions for equations (21,18) depending on the input parameters (**M**, *ρ*, ***τ***), **a**(*t*) and 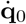. On the other hand, it is in principle also possible that equations (21,18) have no unique solution, which means there is in fact a whole family of strategies with extremal fitness for the same input parameters; this is however not an issue in this study focused on showing that such strategies are possible in the first place. We explore the question of uniqueness of solutions for the EOM in a separate study focused on the deeper mathematical properties of GM, where by reformulating the problem on independent coordinates we can show that solutions must be unique at least for CGMs with a number of reactions that is odd or equal to four. For now, we explore solutions of the EOM with a numerical solver – which by construction picks one solution of the possibly multiple ones – as a tool to study the qualitative properties of the resulting OGTs.

### F. Qualitative properties of the EOM

We solve numerically the EOM of two simple CGMs to illustrate its qualitative properties, in particular the emergent self-oscillation of model L3 caused by: i) an initial condition out of steady-state in a static medium, and ii) a steady-state initial condition on a dynamic medium. We also solve the EOM for model L3 on a periodic medium and for a second, more elaborate model “G5” on a static medium, observing that a more complex quasi-periodic behavior emerges in both cases. We used the daspk function from the R package deSolve [55], designed for solving algebraic differential equations (DAE), see Methods. The EOM (18,21) are first order differential equations and the initial velocities 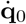 already determine the initial conditions, but for all cases in which we used a “pertubative” non-zero initial velocity 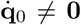, a preliminary check for consistent initial conditions was necessary (see Methods). The following numerical examples are based on highly coarse-grained cell models used here only as a tool to capture qualitative properties of growing cells, not accurate quantitative descriptions. Kinetic parameters were estimated based on measured proteome of *E. coli*, see Methods. For additional CGM examples, see SI Appendix.

#### 1. Model L3 on a static growth medium

We first consider a static medium with initial state 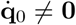 (Fig.2A), which may be seen as representing a cell that was previously at the OGS for its static medium, and after an internal perturbation (e.g. triggered by noise in gene expression) goes into a periodic oscillatory behavior that is manifest in all variables related to its state. The periodic nature of this OGT is more clear in the configuration space **q**, where it forms a closed orbit (see Fig.SI 4A), with frequency *ν ≈* 2.63*h*^−1^ calculated by a fast Fourier transform (FFT) (see Fig. SI 4D). The most important point in this simulation is the clear pattern in the dynamical biomass allocation that indicates the simple “rationale” behind the optimal growth trajectories (OGT) predicted by the equations of optimal motion (EOM): the optimal biomass allocation shifts continuously towards the cell components that are most needed at each point in time, in cycles starting with 1) a peak in transporter protein allocation *ϕ*_ctr_ (in this example with a corresponding peak in growth rate *µ*, since at this time the unique transport reaction can “pump in” more biomass), then 2) a peak accumulation of “carbon” in biomass *b*_C_, which increases the saturation of the following enzymatic reaction, then 3) a peak in the (now more saturated) enzymatic protein allocation *ϕ*_enz_, which then causes 4) a peak in “amino acid” in biomass *b*_A_, corresponding to a peak saturation of the next ribosome reac-tion, and finally 5) a peak in the (now more saturated) ribosome protein allocation *ϕ*_r_, after which another cycle starts. This sequential allocation of proteins to different reactions would require a corresponding sequential allocation of ribosome fractions ***χ*** to produce each of these proteins (bottom of Fig.2A), calculated in these numerical examples by assuming no protein degradation (see Methods). These values ***χ*** may be thought as the fractions of ribosomes bound to each of to the different mRNA (not explicitly modeled in this CGM) encoding the proteins allocated to each reaction, so these simulations indicate also a temporal organization of functionally related transcripts, which has indeed been observed experimentally in yeast [58, 59] and indirectly by promoter activity profiles in the bacterium *E. coli*, a pattern also known as “just-in-time” [60]. Although these studies strongly suggest the corresponding proteome fractions ***ϕ*** must follow the same temporal pattern, there is currently no direct evidence for this due to the technical limitations in measuring dynamical proteomic measurements at genome-scale [61]. A recent study on *E. coli* growth on different steady-state media have found that average concentrations of mRNAs and corresponding protein concentrations at population level are indeed strongly correlated by a “principle of transcriptional predominance” [33], further suggesting protein fractions ***ϕ*** are likely subjected to the same oscillations found in single-cell transcriptomics. Dynamic metabolomic measurements are also currently very limited, but do show a sequential pattern in the oscillations of close related metabolites, for example in the TCA cycle of mammalian cells [62]. Recent single-cell measurements of adenosine triphosphate (ATP) in *E. coli* [63] and ATP, reduced nicotinamide adenine dinucleotide (NADH) and reduced nicotinamide adenine dinucleotide phosphate (NADPH) in yeast [2] showed the concentrations of these essential metabolites oscillate in strong correlation with the cell cycle, further indicating that all key cell functions which involve these metabolites are also bound to be oscillatory at single-cell level.

**FIG. 2.**
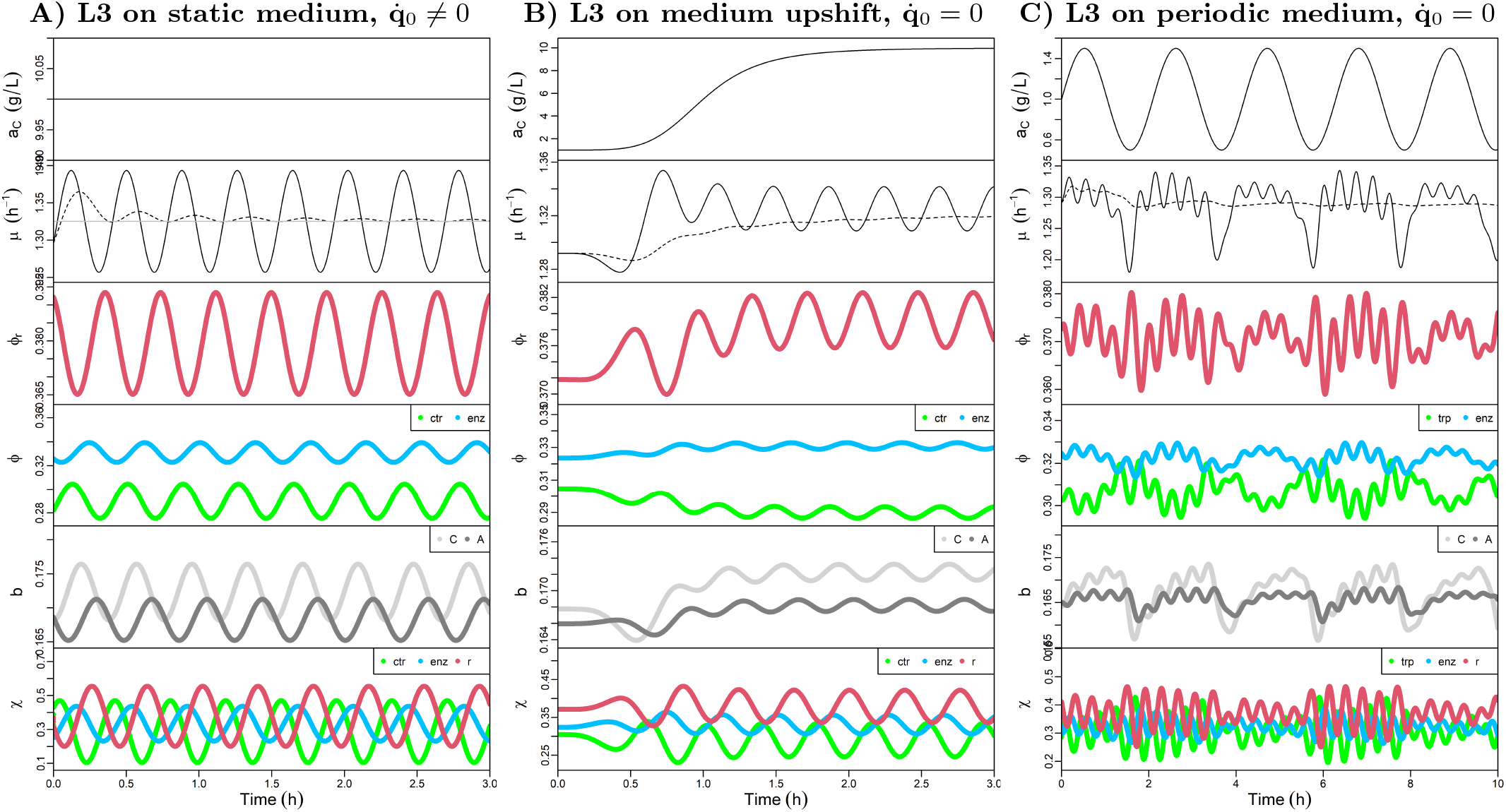
Numerical solutions for the equations of optimal motion (21) of model L3 described in Fig.(1) show emergent oscillatory behavior at different types of growth media. The x-axis in all panels represent time (*h*), while the y-axis represent the medium concentration of “C” *a*_*C*_ (*t*), the (instantaneous) growth rate *µ*(*t*), the ribosome proteome fraction *ϕ*_r_, the remaining proteome fractions ***ϕ***, the biomass fractions **b** (not showing the proteome fraction *b*_p_), and the ribosome fractions ***χ*** allocated to produce each of the proteins. **A)** Simulations for a static growth medium *a*_*C*_ = 10 *g L*^−1^ and initial condition 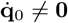. The system presents a self-oscillatory behavior, with the variables in all panels showing the same natural frequency of oscillation *ν* ≈ 2.63 *h*^−1^. The peaks in biomass fraction (each biomass fraction **b** as a whole, and each proteome fraction ***ϕ*** in between) follow the same order as their position in the linear reaction network. The mean growth rate 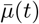 (i.e. the fitness, indicated by the dashed line) converges at longer times to a value Λ about 0.003% higher than *µ*^*\**^ (horizontal gray line). **B)** Simulations for a growth medium upshift defined by a change in the medium concentration according to a sigmoid function *a*_*C*_ (*t*) = 9*t*^5^*/*(1 + *t*^5^) + 1 *gL*^−1^, and initial 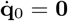. The system initially presents a steady-state behavior corresponding to the OGS for its initial static medium, as expected, until the medium changes provoking a transition phase with an initial decrease in growth rate which has been also observed experimentally for *E. coli* [7, 56], followed by a persistent self-oscillatory behavior again with frequency of *ν* ≈ 2.63 *h*^−1^. **C)** Simulations for a periodic medium where the external concentration changes according to a sine function, and steady-state initial condition 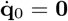. The FFT of *µ* (see Fig.S1) shows the oscillatory behavior can be understood as a combination of the medium oscillation frequency *ν*_m_ *≈* 0.48 *h*^−1^ and the frequency *ν* ≈ 2.63 *h*^−1^, similar to what the model would have if in a static medium *a*_*C*_ = 1 *gL*^−1^ corresponding to the mean *a*_*C*_ (*t*) (not shown). The L3 fitness in this oscillating medium is inferior to the fitness in a static medium with same average concentration *a*_*C*_ = 1 *gL*^−1^, an effect also observed in *E. coli* cells [57].

We identified emergent oscillations in biomass allocation as the underlying strategy of OGTs to get more efficient reactions in order to maximize the average growth rate 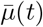 (dashed line in Fig.2A) at every time point *t*. In this simulation, we observe that the average growth rate 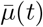 has an initial transition phase but converges at longer time intervals *t → ∞* to a value Λ (the long term growth rate) that is very close to the optimal balanced growth rate *µ*^*\**^ (gray line) at that static medium (in that example, Λ is about 0.003% larger than *µ*^*\**^). This small difference in Λ and *µ*^*\**^ means that this OGT – let’s label it as strategy “A” – starting at a sub-optimal state with initial growth rate *µ*_*A*_(*t* = 0) *< µ*^*\**^ results in a higher long term fitness Λ_*A*_ than any hypothetical “competitor” OGT – let’s label it as strategy “B” – starting (and remaining) at the optimal growth state **q**^*\**^, resulting in a Λ_*B*_ = *µ*^*\**^ (gray line). Interestingly, we observe that the opposite is also true: OGTs starting with higher growth rates *µ*(*t* = 0) *> µ*^*\**^ result in lower long term fitness Λ *< µ*^*\**^ (not shown); by testing various initial states that satisfy the non-negativity of *χ* (not shown), we observed differences in Λ and *µ*^*\**^ that are always small (*<* 0.01%) and would likely be indistinguishable by current experimental methods if these were properties of actual cells. This indicates that for practical purposes, we may interpret OGTs as the growth strategies with the main goal of “trying” to approximate its long term fitness Λ with the corresponding optimal balanced growth rate *µ*^*\**^ of that CGM in that medium. This also means the time average of **q**^*\**^(*t*) converges to a point that is different, but very close to the OGS **q**^*\**^ (in this example about 100 times closer than the initial state **q**_0_, see Fig.S3). This result indicates that while we expect the biomass allocation in individual cells to oscillate, the optimal balanced growth state **q**^*\**^ at a static medium might in some cases still provide a good estimate for the average state of a “population” of OGTs with different initial states; thus, the study of optimal balanced growth might still be relevant to estimate the average behavior of asynchronous cell populations with intrinsic variation in their internal states (here interpreted as different initial conditions for the EOM), or the average state of a synchronized population in time.

#### 2. Model L3 on a medium upshift

In this example we simulate a medium concentration upshift by a sigmoid function, representing the continuous addition of more nutrient in the growth medium. We start the simulation with the steady-state initial condition 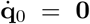, resulting initially in a static OGT while the medium is still not significantly changing. The system then goes through a transition phase while the media shifts to a new steady-state condition, and the OGT finally settles in a oscillatory behavior reminiscent to the example in Fig.2A, with similar frequency *ν ≈* 2.63 *h*^−1^. We observe an “overshoot” in the peak growth rate *µ*(*t*) during the transition phase, which becomes then lower during the following persistent oscillation after the medium shift. That initial “overshoot” followed by a lower peak growth rate has been recently observed in single-cell measurements of *E. coli* going through a medium upshift [7, 56], indicating that the EOM derived here is capable of capturing qualitatively that important oscillatory behavior even in such a simple CGM as the model L3. On Fig.S4 we present other examples of OGT for different types of medium shift, and also observe persistent oscillations after the perturbation introduced by the shifts. Interestingly, we observe that even simpler CGMs with only two reactions (a transport and a ribosome reaction) do not show self-oscillatory OGTs (see Fig. S12), so a minimum of three reactions in CGMs is necessary for self-oscillations to emerge in OGTs. On the other hand, will see later that CGMs with five reactions or more show an even more complex type of self-oscillation. Before that, we explore the optimal response of the model L3 to a periodic medium, as predicted by the EOM.

#### 3. Model L3 on a periodic medium

We simulate now a periodic medium with concentration defined by a sine function, roughly representing cycles such as the feast and famine for organisms in the gut microbiome or the light available during day and night for photosynthetic organisms. We start the simulation with a steady-state initial condition 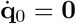, and observe the system quickly goes into an apparent quasiperiodic behavior determined mainly by the oscillation frequency of the medium *ν*_m_ *≈* 0.48 *h*^−1^ and the frequency *ν ≈* 2.63 *h*^−1^ (see Fig.S1F), similar to its own natural oscillation frequency at static medium with the average concentration of this periodic medium *a*_*C*_ = 1*gL*^−1^ (not shown). Interestingly, while the oscillation of the whole system is now defined by two independent frequencies and it is more complex than in the cases illustrated by Fig.2A,B, we still observe the same sequential allocation of biomass to reactants and proteins as before, where the peak investment on each one of these happens in the same sequence as they are found in the linear pathway of model L3. The L3 fitness in this oscillating medium is also reduced when compared to the fitness in a static medium with equivalent average concentration *a*_*C*_ = 1 *gL*^−1^, a type of effect also observed in *E. coli* cells [57].

By testing different frequencies *ν*_m_ and amplitudes for the sine wave defining the medium concentration *a*_*C*_(*t*), we observed that resonance happens in cases the medium oscillations have large amplitudes or a frequency *ν*_m_ too close to the natural frequency *ν* of that CGM in a static medium with similar concentration (see examples in Fig.SI 3), leading to increasingly larger amplitudes in the OGT oscillations until it finally violates some of the non-negativity constraints for **b, *ϕ, χ***. These results could be interpreted as non-physiological states indicating cell death, in an apparent contradiction for an optimal growth strategy. Mathematically, these type of non-physiological solutions would in principle be avoided if the non-negativity constraints (9,10,11) were incorporated as inequality constraints in the 𝒜 functional (20), generalizing our present mathematical framework to a more elaborate, but still relatively typical problem in variational calculus. Since it is not clear whether actual cells could always enforce those inequality constraints, our present non-physiological solutions already capture meaningful predictions and highlight how our optimization here focus on the best possible growth trajectory **q**^*\**^(*t*) for a given (fixed) CGM, but not on optimizing the properties (**M**, *ρ*, ***τ***) themselves defining the CGM; in reality cells growing during longer periods of time eventually adapt via deeper physiological changes (e.g. due to genetic mutations, or horizontal gene transfer) that may be interpreted in our framework as changes in the CGM itself (in the periodic medium example, possibly leading to a different natural oscillation frequency *ν* and thus de-synchronizing with the medium), and thus are not captured by our considerations at shorter time scales (roughly up to hours and days) where the properties (**M**, *ρ*, ***τ***) can be assumed to stay unchanged. When taken at face value, our predicted optimal cell behavior ignoring non-negativity constraints suggest that some cells (as represented by a corresponding CGM) will grow fast but eventually put themselves at a situation of no return and die in certain media, while other cells (represented by a different CGM) will possibly grow slower but consistently in the same media, due to the adaptive properties encoded in their (**M**, *ρ*, ***τ***); that sort of “fitness trap” is in fact observed in nature, e.g. some simpler bacteria growing on a high glucose medium use fermentation as a fast growth strategy but end up excreting too much acids in the medium, eventually causing their own death in what is referred to as “ecological suicide”, while other bacteria have genes encoding extra reactions to cope with their own excreted acids and thus can indefinitely grow on the same medium [64].

We now explore how quasi-periodic behavior emerges even in static media for a CGM with five reactions.

#### 4. Model G5: five reaction model including replication and transcription

We now consider a less trivial CGM with 5 reactions, representing a general cell growth model with the “central dogma” of biology including replication (via DNA polymerase “DNAp”) and transcription (via RNA polymerase “RNAp”). In that model we consider a more general kinetic rate law including activation (see SI Appendix …), so DNA here needs to be produced in order to activate its own production (i.e. DNA replication) and the production of RNA (i.e. transcription), which in turn needs to be produced in order to activate the ribosome reaction (i.e. translation), a simplified way to account for how different types of RNA need to be produced to allow translation, including the rRNA composing part of the ribosome itself. Fig.(3)A presents the scheme of the model and Fig.(3)B its matrix **M**. This model is also more realistic by including branched pathways instead of the simple linear pathway found on model L3, exemplifying how our results are general and apply to arbitrary reaction network structures.

**FIG. 3.**
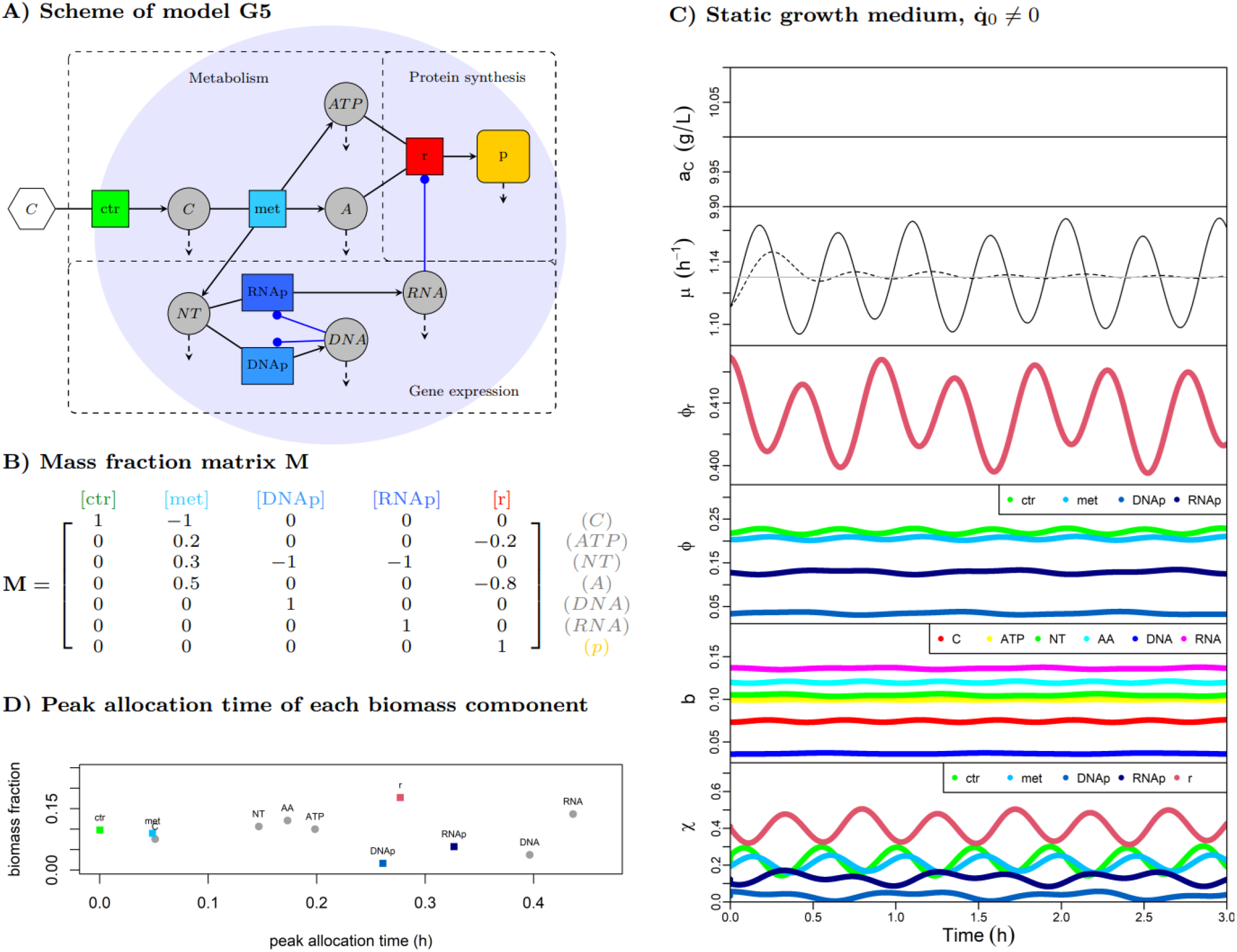
**Model G5** with 5 reactions (black arrows) including a coarse-grained representation of the “‘central dogma” of biology with DNA replication, RNA transcription, and protein translation (a more accurate name for the “ribosome reaction”). The net production of each reactant due to dilution by growth and concentration changes in indicated by the dashed arrows. Here we can identify in the “non-protein synthesis” two subnetworks representing metabolism and gene expression. **A)** Representation of the reaction network structure with branched pathways instead of the simple linear pathway of model L3, and blue arrows indicating activation; here DNA and RNA are produced from “nucleotides” (NT) but not consumed as substrates by any reaction. However, DNA production is necessary to activate its own production and the production of RNA (blue arrows), and RNA production is necessary to activate the ribosome reaction producing proteins (blue arrow), which here also require ATP produced by the metabolic reaction met. **B)** The model G5 is completely defined mathematically by the mass fraction matrix **M**, the density *ρ* = 340 *gL*^−1^, and turnover times ***τ*** following a kinetic rate law with activation (see SI Appendix, and kinetic parameters in Fig.S7). **C)** The results for a *T* = 3 *h* simulation based on the numerical solution of the corresponding EOM at a static medium *a*_*C*_ = 10 *gL*^−1^ but non steady-state initial condition 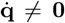. The emergent oscillations are here quasi-periodic, with two fundamental frequencies as determined by a FFT (see Fig.SI8). Again, the dashed line represents the average growth rate at each point in time, and the gray line the growth rate at the OGS. **D)** The peak allocation time for each biomass component in this model at the first simulated growth cycle of about 0.5 *h*, showing a trend towards the sequential allocation observed in L3, but which is in fact more complicated specially within the components involved in the “gene expression” subnetwork.

We then solve numerically the corresponding EOM (21,18) for the model G5 in a static medium with non steady-state initial conditions 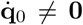, which results in an oscillatory OGT (Fig.3C) as for the simpler model L3, with the main difference that now the oscillation is apparently quasi-periodic with two fundamental frequencies as indicated by a FFT of *µ*(*t*) (see Fig.S1F). The reaction network in the model G5 is not a linear pathway, but we can still identify the same sequential biomass allocation strategy in its backbone reactions resembling model L3 (Fig.3D): transport proteins peak first, followed by C, the metabolic proteins “met”, then the intermediate reactants between met and r reactions (A, NT, ATP) and finally ribosomal proteins r. We observe DNA and DNAp peak around the end of the cycle, while RNA peaks close to the ribosome when it is most required to “activate” the translation. Here and in other less trivial CGMs with 5 or more reactions (see models L5 and G9 in SI Appendix), the just-in-time strategy can still be observed as an overall trend in the optimal dynamical resource allocation across different pathways, while more complex patterns emerge within the pathways themselves, as have been also observed for amino acid biosynthesis pathways in *E. coli* [47].

The observation of this hypothetical quasi-periodic behavior in actual cells poses a difficult challenge for experimental methods, but a recent study with detailed measurements have indeed found evidence for a quasiperiodic ATP oscillation in single *E. coli* cells, with one major frequency correlated with the cell cycle and second faster one [63]. Since ATP is a central molecule for life, it is possible that such quasi-periodic behavior will be found to be wide-spread across organisms and cell com-ponents once experimental methods advance and become sensitive enough to detect it.

We tested the numerical solutions for the EOM of various CGM and observed that the quasi-periodic behavior apparently emerges depending solely on the number of reactions in a CGM, as we discuss next.

### G. Emergence of periodic and quasi-periodic self-oscillations

In addition to models L3 and G5, we tested the numerical solutions for the EOM of other CGMs on static media with non steady-state initial conditions 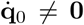, and found the type of self-oscillation in their corresponding OGT to be determined exclusively by the number of reactions in the CGM, as summarized on Table I. The simplest possible CGMs with only two reactions (transport and ribosome) do not show any self-oscillation in static media, and only damped oscillations in dynamical media (see Fig.S16). All CGMs with three and four reactions (see three examples in SI Appendix) show periodic self-oscillation in static media, indicated by a close orbit in the configuration space **q**(*t*) and a single frequency in the FFT of the growth rate *µ*(*t*). All CGMs with 5 or more reactions (see three examples in SI Appendix) show apparent quasi-periodic OGTs in static media, as indicated by their trajectory in the configuration space **q**(*t*) and more than one fundamental frequency in the FFT of *µ*(*t*).

**TABLE I.** General pattern observed in the type of self-oscillation in numerical solutions of the EOM at static growth media for CGMs with different number of reactions, as indicated by a FFT of the growth rate at longer simulations with *T* = 100 *h*. For the simplest type of CGMs with only two reactions, we observe no self-oscillation. For three reaction models (such as L3) and 4 reaction models (see examples in the SI Appendix), we observe periodic oscillations for all CGM tested (independent of network structure), with closed orbits in the configuration space **q**(*t*). For all CGMs with five or more reactions tested (including a linear pathway model L5), we observe numerically (see SI Appendix for examples) only apparent quasi-periodic behavior (independent of network structure), as indicated by a FFT of *µ*(*t*).

| #(reactions) | Self-oscillation in static media | #(frequencies) |
| --- | --- | --- |
| 2 | not existent | 0 |
| 3 | periodic | 1 |
| 4 | periodic | 1 |
| 5+ | quasi-periodic | 2+ |

We also observe in numerical solutions that the self-oscillation frequencies are largely independent of the initial conditions 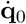, unless at extreme initial values 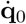 corresponding to non-physiological states with negative ribosome fractions ***χ*** (see examples in Fig.S4). This frequency *ν* for each CGM however strongly depends on the static medium considered, and by consequence a relationship emerges between *ν* and the corresponding long term growth rate Λ across different static media.

### H. Emergence of “growth laws” across different static media

We solved the EOM numerically for different CGMs in different static media, and observed that for each CGM an quasi-linear relationship emerges between its oscillation frequency *ν* (in case of 5+ reaction CGMs, its main frequency as determined by a FFT of *µ*(*t*)) and its long term growth rate Λ across the different media (Fig.4A). In all cases, we observe that a linear regression at the tested regimes of intermediate to high Λ points to an apparent non-zero frequency “offset” *ν*_0_ at the zero growth limit Λ *→* 0, a qualitative behavior that has been observed experimentally in different strains of yeast [65] and emerges here exclusively from the first principles of fitness optimization and basic constraints on cell growth (6-8), instead of being itself assumed as a phenomenological law between these cell properties. Since we made no reference to the cell cycle in our assumed constraints on cell growth (6-11), the oscillations predicted here are more naturally identified with what is referred as’metabolic oscillations in cells, in contrast to the cell cycle oscillations related to the specific cell components produced at the cell division times. Nevertheless, various studies indicate that both types of oscillations are coupled in microbes [2, 63, 66], in a one-to-one fashion or in a “one-to-many” fashion when single cells commit to the cell division cycle only at a multiple number of metabolic cycles [65]. This necessary coupling between metabolic and cell cycle oscillations due to physiology can be seen as an extra biological quantization rule that constrains the timing of cell division, which we do not account for here but must integrate in the future developments of the theory.

**FIG. 4.**
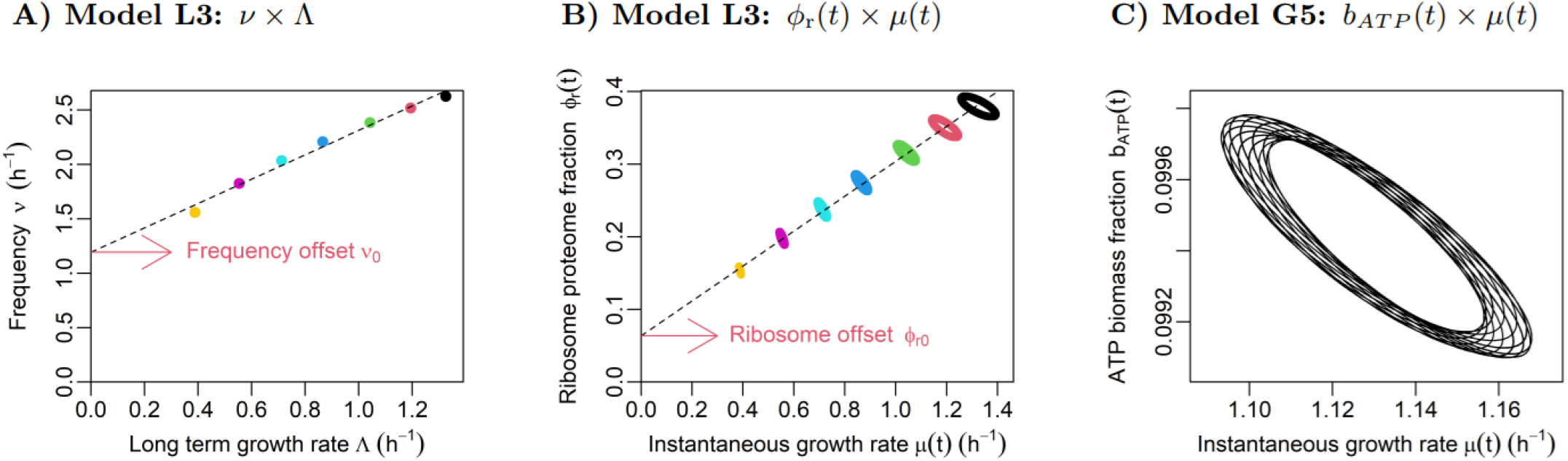
Emergent relationships between optimal cell properties across different static media (different colors) for models L3 and G5 (see Fig.S20 with similar qualitative results for other CGMs). **A)** We observe the natural frequency *ν* relates in a quasi-linear way with the long term growth rate Λ of model L3 across different static media, with linear regression (*r*^2^ = 0.98) resulting in a frequency “offset” *ν*_0_ ≠ 0, which is consistent with the experimental observations for the metabolic oscillations in some yeast strains [65]. **B)** Solutions for different static growth media (each color corresponding to the same medium as in A) resulting in periodic ribosome proteome fraction *ϕ*_r_(*t*) and instantaneous growth rate *µ*(*t*), with each “ring” corresponding to a anti-clockwise motion in that plane; this results in a negative correlation between these variables in each medium, while the average values in each medium (not shown) correlate positively (dashed line) across the media, a type of phenomenon known as Simpson’s paradox. The mean values 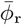 and 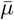(= Λ at infinite time) follow an quasi-linear “growth law” (*r*^2^ = 0.99) with a non-zero “offset” 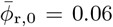, in qualitative agreement with the experimental results for various microbes [26, 27, 46]. All simulations here consider the maximal possible initial value for the ribosome generalized velocity such that the non-negativity of ***χ*** is not violated, resulting in lower oscillation amplitudes at lower 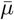 (see Methods). We observe the same basic qualitative behavior in all other CGMs tested, with closed orbits (as in this example) for CGMs with three or four reactions, and nonperiodic orbits for CGMs with more than four reactions (pattern in Table 1), see examples in Fig.S20. **C)** The relationship between ATP in biomass *b*_*ATP*_ (*t*) and instantaneous growth rate *µ*(*t*) predicted by the EOM (21,18) for model G5 at same initial condition as Fig.(3C) in and *T* = 10 *h*, which is consistent with the reported anti-correlation between these properties in single *E. coli* cells [63].

We also inspected in our simulations the most commonly described growth law relating the (population average) ribosome proteome fraction 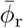 and the long term growth rate Λ across different static media by a linear relationship 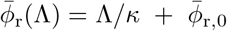 (with *κ* and 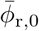 being positive constants), as it has been many times observed experimentally and used as a modeling input in various descriptive cell models [26]. We found that our dynamical simulations at long time *T* = 100 *h* predict an average ribosome proteome fraction 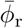 and average growth rate 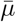 (used now on as a proxi for Λ, the average growth rate at infinite time) that indeed follow a quasi-linear relationship at intermediate to high growth rates (see Fig.4B for the results of model L3, and Fig.S20 for the same qualitative result using other CGMs), with a non-zero ribosome offset 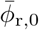 at the zero growth limit as described by the well established growth law [26] which has recently been show to be valid as population averages, but not in single *E. coli* cells [8]. Interestingly, when we inspect our actual dynamical results, not the averages – which could be interpreted as predictions for single-cells, instead of how the averages mentioned before would correspond to averages for populations of de-synchronized cells at different points of the growth cycle – they follow in each medium a type of dynamic growth law anti-correlating *ϕ*_r_(*t*) with *µ*(*t*) (Fig.4B), which to our knowledge has not yet been directly tested by experiments and thus represents an independent new prediction that could be tested to validate the assumptions used for our mathematical derivations. Again, this qualitative prediction is based exclusively on first principles, and in case of being confirmed by experiments, would help guiding the study of cell growth, e.g. as an additional ingredient in mechanistic models of single growing cells. An anti-correlation has however been recently reported between dynamic levels of ATP in biomass *b*_*AT P*_ (*t*) and instantaneous growth rate *µ*(*t*) in single *E. coli* cells [63], which we also observe in our simulations of model G5 (Fig.4C) – where ATP is directly connected to the ribosome reaction, capturing the fact that most ATP in cells is consumed by translation and influenced by its dynamics [67] – further indicating these simulations of highly coarse-grained CGMs are indeed capturing qualitatively some important properties of growing cells.

## III. DISCUSSION

In this work we introduced Growth Mechanics (GM) as a general mathematical framework to formalize and solve the optimization problem underlying the natural selection of growing cells: for a given dynamic growth medium, how should a cell optimally allocate its resources in time, so that its average growth rate in a certain time interval (i.e. its fitness) is maximal, under the constraints of mass conservation (including dilution of all components by growth and their concentration changes), arbitrary (in general nonlinear) kinetic rate laws, and limited cell density. We formalized this mathematical problem by showing i) how all the relevant information about the system of interest can be encoded in a triple (**M**, *ρ*, ***τ***) defining a “cell growth model” (CGM) and a function **a**(*t*) defining the relevant concentrations in the growth medium, and ii) how the description of the internal system state in each point in time can be simplified by a minimal set of “generalized coordinates” **q**(*t*) and “generalized velocities” 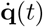, in close analogy with the mathematical framework of classical mechanics. That key mathematical simplification in the description of growth states leads to the formalization of the optimal dynamical growth problem in terms of a “principle of extremal fitness” (20), analogous to how the principle of extremal action defines physical systems using the formalism of variational calculus [68]. From that point, we follow the same general procedure employed in fundamental physical theories: we solve the corresponding Euler-Lagrange equations and find here the “equations of optimal motion” (EOM) (21,18), a system of differential equations determining the “optimal growth trajectory” (OGT), the optimal system dynamics for a given initial condition. We then showed with numerical examples for simple CGMs how oscillations emerge in OGTs, as long as both cell state and medium are not initially at steady-state. We also observed how the predicted optimal behavior across different static media results in the emergence of the known “growth law” between the average ribosome protein allocation and growth rate in each growth medium, and a new type of “dynamical growth law” between these properties in time, which have not yet been tested by experiments. The identification of such oscillations in experimental data – specially at small amplitudes, high frequency, and in single-cells – is challenging due to the confounding effect of noise; in fact, most of the current studies on single-cells at static media tend to interpret dynamic fluctuations as noise only, while the periodicity of fluctuations is usually acknowledged for microbes which synchronize and show oscillations more evidently at population level, such as in yeast [2, 6, 59, 65]. The treatment of noise in theoretical studies could also in some cases potentially mask the presence of persistent oscillations in experimental data, for example when averaging the state of multiple cells in a de-synchronized cell population [40] or using moving averages to smooth the noise in single-cell data [7].

### A. GM as a unifying framework to model growing cells

Our mathematical framework can be seen as a generalization of previous optimization-based approaches, which in order to simplify the difficult mathematical problem underlying optimal cell growth and reduce the number of necessary kinetic parameters used as inputs (which are still very scarce in literature [69]), account in each case only partially for the constraints on growth described here. Often the mass conservation constraint (6) is approximated by ignoring either i) changes in concentrations (i.e. balanced growth regime), ii) dilution by growth, or iii) both. These approximations are sometimes i) taken for all cell components [50], ii) taken for all non-protein components [29, 49] and usually referred as “quasi-steady-state” approximation, or iii) accounted only approximately for a subset of proteins and non-proteins composing the most “important” part of the cell biomass, as judged by experts in each particular case [52]. These approximations are useful in some contexts by resulting in greatly simplified optimization problems that depend on fewer input parameters and can be solved efficiently by numerical methods, thus providing a practical way to estimate cell properties with relative success, even at genome-scale in case of linear models [19, 70]. These conceptual simplifications on what cells need to produce however break the defining feature of true self-replication, and thus limit a deeper understanding of the most fundamental trade-offs in cell economics by these theoretical approaches. Theoretical results on cell growth and metabolism are also often restricted by the assumption of specific kinetic rate laws, including the simple massaction kinetics [50] that cannot capture the saturation of protein catalysts and thus miss important trade-offs in cell economics such as the balance between enzyme and substrates at optimal states [30]; while our numerical results here consider CGMs with the simplest type of nonlinear rate laws (Michaelis-Menten and Michaelis-Menten with activation) for the purpose of illustration, the EOM derived here are analytical results that apply to any kinetic rate law via the given functions ***τ***. Most theoretical works also assume a constant protein in cell biomass *b*_p_ in order to account for the limitation in this important cell resource, while in fact different experimental studies indicate total protein in cells may change significantly for different organisms and different growth media [40]. Here we do not impose a fixed *b*_p_ as an input in the CGMs, but instead have it as an output in the OGTs predicted by the EOM; in our simulations with different CGMs we observe an overall decrease in the fraction of protein in biomass *b*_p_ at higher growth rates (Fig.S21), consistent with some experimental results for the bacterium *E. coli* [40], from which proteomic data we based our estimations of kinetic parameters (Methods).

Our results also generalize previous theoretical studies on optimal dynamical cell growth by solving the optimization problem with the average growth rate (20) as objective function, while previous approaches simplify the objective function with phenomenological assumptions including i) previous knowledge of the final biomass composition **b** cells want to achieve at finite time [71, 72], ii) imposing a limited discrete number of possible “switches” in the system state [45, 73], iii) optimizing only for the long term growth rate Λ at infinite time [74], and more often iv) optimizing the instantaneous growth rate *µ*(*t*) at each point in time, as a proxi for the actual fitness measure 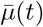 (in linear models, in fact the yield optimization for a given biomass **b** is used itself as a proxi for *µ*(*t*) [75]). Another approach on modeling growing cells ignores the optimization of an objective function, but assume instead that the long-term dynamics must satisfy some ergodic condition [10]. Interestingly, while our definition of CGM fits in the broader concept of scalable reaction networks (SRNs) defined by Lin et al. [10] (which allow for multiple protein synthesis reactions), their ergodic assumption on the longterm network dynamics is not strictly true for OGTs, but holds as a good approximation for the simple CGMs tested here; in particular, that assumption implies that the long term growth rate Λ would not depend on the initial conditions, which we have seen it does to a small extent for the OGTs found here.

### B. On modeling cell growth via optimization principles

The study of natural phenomena via an optimization perspective goes back to the natural philosophy of Aristotle, who summarized this guiding principle by stating “If one way be better than another, that you may be sure is nature’s way.” [76]. In modern science, the quantitative use of an optimization principle to explain physical phenomena begins with Fermat’s principle on how light always follows a path that minimizes the time to travel between two points [68]. This general idea later evolved as a unifying framework for all physical systems via the principle of extremal action, a quantity defined as the integral of a Lagrangian function in time for each specific system [68]. While the practical success of this approach in physics is evident, the fundamental meaning of both concepts of action and Lagrangian are still debated [77]. On the other hand, modern biology has a foundation on Darwin’s principle of evolution by natural selection and the consequent concept of survival of the fittest, well understood qualitative principles that intrinsically involve an optimization view of natural biological phenomena [76], but with no general unifying framework for quantitative predictions. Here we showed how, in the particular case of growing cells with no strong influence on their growth medium state (so the function **a**(*t*) is a given, independent input), we can formalize a general quantitative principle of extremal fitness. As for the principle of extremal action in physics, the resulting optimal dynamical behavior is defined by a set of differential equations and some given initial conditions only; this means that the optimal solution **q**^*\**^(*t*) at each point *t* in time depends only on the initial condition and corresponding instantaneous medium state 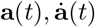, but it is independent on the future states of the medium. That mathematical result may seem counter-intuitive, in the same way that its analogous problem in physics has also sparked a continuous debate on how an effect (the optimal trajectory) can apparently precedes its own cause (the extremization of a functional at a final time *T*), known as the teleological problem in metaphysics [77]. As for the working physicist, this teleological concerns do not however limit the potential use of this type of variational principle to understand the physical systems we are interested, specially considering i) here fitness is a more concrete concept than action, with its assumed optimality grounded on the well established theory of evolution and ii) the optimal behavior predicted here by the EOM are evidently not supposed to be exactly followed by actual cells (as equations of motion are usually supposed to be exactly followed in physics), and should be understood only as the limit case guiding the study of growing cells. We emphasize that the simple L3 and G5 models are presented here only to illustrate the qualitative behavior predicted by the EOM, and are not fine tuned for accurate quantitative predictions. In practice, the optimization approach to model cell growth is already the most prevalent, including in the simplest and most common frameworks to model steady-states (flux balance analysis (FBA) [18]), and dynamical cell growth (dynamic FBA (dFBA) [75]).

### C. The adaptive properties encoded in CGMs

In this work we focused on finding the hypothetical optimal growth strategy in time **q**^*\**^(*t*) for a given (possibly dynamical) growth media **a**(*t*), and CGM defined by the properties (**M**, *ρ*, ***τ***) that are assumed to be fixed in time. This means i) different CGMs may in principle respond differently to the same given medium, as predicted by the EOM, and ii) our assumptions only hold as long as as the properties defining the given CGM remain themselves unchanged in time. If it is possible that after some long enough period of time a particular cell becomes perfectly adapted to its (somehow predictable) growth medium, this would mean in our framework that all the information about the medium become built in the CGM itself, through its structure encoded by the matrix **M**, properties of kinetic functions ***τ*** and density *ρ*. Since each different model has its own optimal response to a growth medium (as determined by the EOM), we might see in this perspective the information encoded in a particular model (**M**, *ρ*, ***τ***) as an “algorithm” to thrive (to some level of success, not perfectly necessarily) in a certain kind of growth medium **a**(*t*). Following this line of though, we can also think of the natural selection of cells with changes in the properties captured by (**M**, *ρ*, ***τ***) (due to e.g. genetic or epigenetic changes) at longer time scales as a way to better react to some fundamental change their respective growth medium (e.g. a significant change in the frequency of a periodic medium). We can thus picture the general evolutionary path of unicellular organisms as a compound process where the EOM would be relevant for the (optimal) dynamics at shorter time scales, but changes in the properties of the model (**M**, *ρ*, ***τ***) must also be taken into account at larger time scales. A recent study has shown evidence that distantly related bacteria (each adapted to a different growth medium) show a surprisingly conserved proteome allocation ***ϕ***, while their kinetic properties (most importantly, the ribosome turnover time *τ*_r_) differ significantly [78], possibly due to different saturation levels but also changes in kinetic properties that could happen at long evolutionary time scales.

### D. Relationships with previous theoretical results on cell growth

Previous theoretical studies have indicated that a sequential “just-in-time” pattern in protein allocation along the metabolic network optimizes proxies of cell fitness, under the assumption that the system must converge to a steady-state when the growth medium also shifts to a steady-state [60, 73, 79]. A discrete sequential protein allocation in time has also been shown to generally increase fitness in a linear optimization-based model of yeast cells in steady-state medium [6]. In a recent publication, one of the present authors demonstrated that small, persistent oscillations around the optimal steady state in kinetic models of enzymatic pathways can enhance efficiency – defined as the average flux per average enzyme activity – thus challenging the prevailing assumption that optimal metabolic states in steady media must themselves be steady states [51]. Here, we generalize this theoretical result by showing that continuous oscillations naturally emerge in the global biomass allocation of cell growth models (CGMs), which include the necessary synthesis of proteins to catalyze all reactions, and are derived entirely from the first principles of fitness optimization (quantified by average growth rate), mass conservation (including dilution by growth and concentration changes of all components), kinetic rate laws, and constant cell density. While the function of changing cell states in fluctuating growth media have been understood previously as a way to improve fitness by general theoretical studies [57, 80] and for specific models of particular organisms in transient or stead-state media [5, 7, 50, 81], our results provide a unifying rationale to explain the observed oscillation of microbes as a way to improve their fitness in general, including by self-oscillation on steadystate media. This general rationale also helps explaining the wide-spread observation of simple linear relationships between cell properties across different media, including the ubiquitous growth laws relating ribosome proteome allocation and growth rate [26].

Our results also differ from many of the previous theoretical studies on optimal cell resource allocation that are built as an optimal control problem [45, 74, 79], where some cell properties are treated as control variables (e.g. the ribosome fractions ***χ*** allocated to produce different proteins). Our modeling framework implements instead a holistic view of the growing cell, which does not differentiate between which cell components have the role of controlling others; the whole internal cell state at each point in time 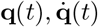 is determined as consequence of the fitness optimization at a given medium state 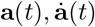. In this sense, the GM framework does not impose any intrinsic hierarchy on which cell properties are controlling others. A possible extension of the present framework could however consider the medium concentrations **a**(*t*) as control variables, a scenario of great importance e.g. in industrial cultures.

### E. Limitations in the current framework and possible generalizations

In this study we focused on a minimum set of mathematical constraints capable of explaining the main properties of cell growth. While this simplifying approach helped finding quantitative principles relevant for growing cells in the most general sense, some additions would help capturing more specific features of cell growth. One is the generalization to models with arbitrary matrices **M** including linearly dependent columns, important to understand how certain reactions become active or not in particular growth conditions [16]. Another important generalization is the addition of more constraints, e.g. the degradation of macromolecules which become a significant feature for slow growing cells, and the inequalities (9-11) enforcing non-negative fractions of protein ***ϕ***, biomass **b**, and ribosome ***χ*** allocation, which must be incorporated to fully understand how and when these become limiting to optimal growth. In all the simulations presented here, these non-negativity inequalities were satisfied and did not constrain the optimal states, but inequality constraints in but that would not be the case for larger perturbations in the initial conditions or faster media changes, in particular for ***χ*** (see Fig.SI4) which showed in all cases much larger proportional variations than the other model properties and thus could not be used as primary coordinates to model faster medium transitions without becoming negative [40]. The large variations in ***χ*** observed here indicate that other factors besides ribosome allocation and dilution by growth may be necessary for better modeling protein allocation in faster changing media (when nonnegative ***χ*** become a limiting factor), such as protein degradation and post-translation modifications. Finally, a more realistic description of growing cells must account for the fitness of cell populations as a whole [48, 80] and how the their own activity influences the growth medium via exchanged reactants. The present study is applicable to regimes of low influence of the cell activity on its growth medium dynamical state **a**(*t*), which is assumed to be known. While such conditions can be achieved in controlled experiments, the realistic modeling of large cell populations in nature requires the coupling of both medium-to-cell and cell-to medium influences, e.g. in fasting-famine cycles [39] and synchronous population growth [59]. The interaction between cell subpopulations via their shared growth media is also an important factor in natural populations, and must be accounted for to fully understand their dynamics and emergent behavior as a whole [8]. We further explore some of these generalizations in a separate theoretical study, where we also clarify the analytical basis for the pattern observed in Table (I) and show how the mathematical structure of GM is closely related to that of classical field theories.

While our optimization focused approach does not require the full knowledge of specific mechanisms implementing the cell states – in particular, the oscillatory resource allocation discussed here – to derive general principles for growing cells, a recent study has shown that the position of gene families in the bacterium *E. coli* is biased toward specific chromosomal positions [82], indicating gene expression could itself be to some extent an ordered process facilitating cells to get closer to the optimal oscillatory growth strategies. We finally suggest that the development of highly curated models in the GM framework would built on the knowledge of existing descriptive models of specific microbes (e.g. [5, 46, 47]), in a complementary process that could progressively bridge the gap between these different modeling approaches.

## IV. METHODS

### A. Models parametrization

We determined rough estimations for kinetic parameters based on the following procedure using i) the singular value decomposition (SVD) of each matrix **M**, ii) an assumed typical saturation level of reactions, iii) given density *ρ*, and iv) an initial estimation for the proteome allocation ***ϕ*** and growth rate *µ* (described more generally and in more detail in a separate article in preparation):

- The singular value decomposition (SVD) of each matrix **M**

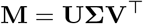

provided in each case a natural choice for positive vectors **q, b** satisfying both equations **Mq** = **b** (12) and **s** · **q** = 1 (18); for each matrix with *n* columns (= number of reactions), the SVD provides in *n* pairs (**q, b**) satisfying equations (12,18), one pair possibly strictly positive. We find these by taking the columns of **V** and scaling each in order to construct vectors **q** satisfying the continuity constraint (18), with the corresponding **b** given by (12). For the L3 model, we have one positive pair with **q** = (1, 0.801, 0.445) and **b** = (0.198, 0.356, 0.445). For the model G5, we have one positive pair with **q** = (1, 0.902, 0.093, 0.093, 0.249) and **b** = (0.097, 0.105, 0.084, 0.276, 0.093, 0.093, 0.249).
- From the estimated **b** and given density *ρ* of each model, we estimate one Michaelis constant *K*_*m*_ = *ρ b*_*m*_*/K*_ratio_ for each internal reactant *m* excluding the total protein, i.e. each internal substrate concentration *ρ b*_*m*_ is *K*_ratio_ times its corresponding *K*_*m*_ (same for all reactions it is involved). We chose *K*_ratio_ = 3, matching average experimental values in *E. coli* [30, 34]. Other similar values for *K*_ratio_ can be easily tested using the provided R codes discussed later, and do not change the qualitative results present here.
- From the kinetic equation (5), and the equation (15) assuming balanced growth 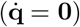, we have

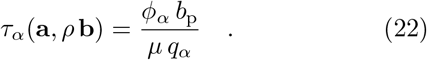

Assuming *τ*_*α*_ follows the Michaelis-Menten kinetics (see SI Appendix) with full saturation of external medium reactants and *K*_*m*_ = *ρ b*_*m*_*/K*_ratio_ as before, we have

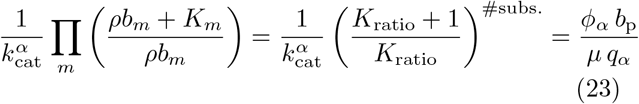

where 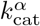 is the turnover number of reaction *α* and #subs. the number of substrates in that reaction. We now solve for 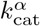 *K*_ratio_ = 3

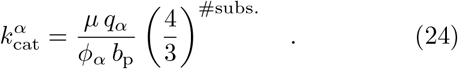

We now estimate each 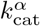 based on the previously calculated *q*_*α*_ and *b*_p_ and experimental proteome ***ϕ*** and growth rate *µ* measurements for *E. coli* culture at balanced growth. For model L3, we estimate ***ϕ*** = (0.35, 0.40, 0.25), where *ϕ*_r_ = 0.25 comes from the whole translational machinery measured in Ref. [47] at growth rate around *µ* = 1 *h*^−1^, the transporter *ϕ*_ctr_ = 0.35 comes from the estimated fraction of non-cytosolic protein in Ref. [31], which leaves *ϕ*_enz_ = 0.40 for the enzymatic reaction. For model G5, we estimate ***ϕ*** = (0.35, 0.30, 0.1, 0.1, 0.15) again at *µ* = 1 *h*^−1^, where *ϕ*_r_ = 0.15 represents the ribosome proteome fraction as measured in Ref. [47].

All estimated kinetic parameters are then rounded to the next integer for the simplicity of presentation and to emphasize these are only rough estimations. For model G5 we also estimate activation constants 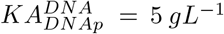 and 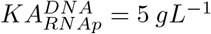 for the DNA activation of reactions *DNAp* and *RNAp*, and 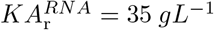 for the RNA activation or the ribosome reaction, equivalent to half of the DNA and RNA concentrations in *E. coli* [83] for the assumed dry weight density *ρ* = 340 *gL*^−1^.

### B. Numerical solution for the equations of optimal motion

The equations of optimal motion (21,18) compose a system of differential-algebraic equations (DAE), which means some choices of initial conditions may not be consistent (another example being the classical pendulum, for which some initial velocities are not tangent to the circle of possible trajectories). The consistency of initial conditions is not an issue in the simulations with 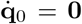 since in those cases the corresponding EOM at *t* = 0 reduces to a simpler system of algebraic equations. In the cases we want to introduce some “perturbation” on the initial conditions (e.g. Figs 2A and Fig.3) we circumvent this consistency problem numerically by first taking the numerical solution for a given initial condition (for model L3 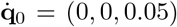 and for model 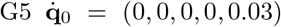) and use its value at the second time point (a consistent condition according to the numerical solver) as the actual initial condition for the EOM we want to solve; for model L3 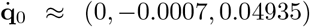 and for model G5 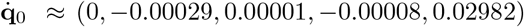. We used the daspk function from the R package deSolve [55], designed for solving algebraic differential equations (DAE), and implemented a general R script for the simulation of small CGMs based on simple inputs defined by open source spreadsheet files (assuming a general type of kinetic rate law [84]). The daspk function accepts systems of equations with the same number of variables as equations, while the EOM (21,18) contains one more equation than its number of variables; we solved that numerically by adding the continuity equation (18) to all equations (21) with a scaling factor of 10^8^ to guarantee a tight conservation of mass. The daspk function also required an initial estimation of **q**_0_, for which we used in all cases the solution of the EOM satisfying initial steady-state 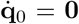. The models L3, G5 with respective media conditions (given functions of time) are defined using open source spreadsheets and can be easily edited, as well as other examples of CGMs given in the Appendix. The simulations are for a given time *T*, time intervals Δ*t*, and can be done by automatically estimating kinetic parameters for a given *K*_ratio_ using the procedure explained before, or by providing parameters in the spreadsheet files. We encourage the use of this code to explore different parameters on the given CGMs, as well as creating other small models and media conditions (defined as functions of time) that can also be simulated using the same code.

### C. Proteome fractions *ϕ*

From equations (5,15,17) we have that each proteome fraction *ϕ*_*α*_ is uniquely determined by 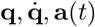 as

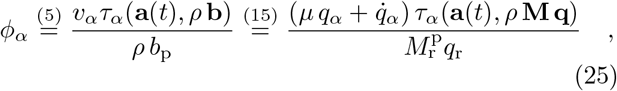

where *µ* is uniquely determined by 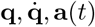 via equation (17).

### D. Ribosome fractions *χ*

Each *χ*_*α*_ quantifies the fraction of ribosomes allocated to produce protein *α*. Thus, for each *α*, mass conservation dictates that for a total protein production flux 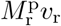 and protein *α* concentration *ρ b*_p_*ϕ*_*α*_

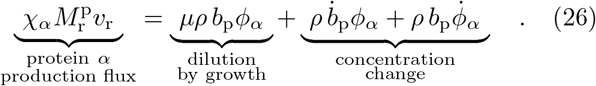

Solving for *χ*_*α*_ and simplifying (see SI Appendix), we get

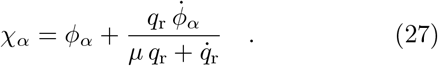

The values of each *χ*_*α*_ in the numeric simulations are then calculated by using equation (27) with the values of *µ, ϕ*_*α*_ calculated from the solutions 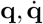, and 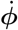 calculated numerically from the changes in ***ϕ***.

## E. Software Availability

All results presented in this work can be reproduced with the R scripts deposited in github at https://github.com/HDourado/Growth_Mechanics.

## F. Awknowledgements

We thank Antonio Rigueiro for guidance on numerical solvers for differential equations, Rainer Machné for insightful discussions on the nature of metabolic oscillations, and Diana Széliová for helpful comments on the manuscript. This work was supported by the German Research Foundation through grant CRC 1310 and under Germany’s Excellence Strategy, grant EXC 2048/1 (Project ID: 390686111).

