## Supplementary Appendix for "Growth Mechanics and the emergence of metabolic oscillations in growing cells"

### Supplementary Appendix for
### Growth Mechanics and the emergence of metabolic oscillations in growing cells

**Contents**


**On notation** 1

**Definitions** 1
Column sum vector $\mathbf{s}$ 1
Kinetic rate laws 2

**Derivations** 2
The continuity equation 2
Ribosome fractions $\boldsymbol{\chi}$ 2
The equations of optimal motion (EOM) 3

**SI Figures** 5
Figures relating to GCMs L3 and G5 in the main text 5
More GCM examples: L2, L4, Y4, L5, G9. 13
Peak allocation time 23
Growth laws for frequency and ribosome allocation 24
Growth law for protein allocation 26

**References** 27


#### ON NOTATION

In the following derivations, it will be useful to represent entries of column vectors by upper indices, and entries in row vectors by lower indices, so we can indicate vector and matrix multiplication with the Einstein summation convention, where each repeated lower and upper index indicates summation. We use indices $\alpha, \beta$ for reactions, with r indicating the ribosome reaction. We use the index $\iota$ for internal reactants, with p indicating the proteome pool, corresponding with the last row $M_\text{p}$ of $\mathbf{M}$, and the last entry $b_\text{p}$ on the vector $\mathbf{b}$ indicating the proteome mass fraction in the biomass. We use the index $\epsilon$ for the external concentrations $a_\epsilon$ defining the growth medium. We use the standard convention that the Greek indices start at zero, so e.g. in model L3 we have the coordinates $q^\alpha = (q^0, q^1, q^2)$, where $\text{r} = 2$.

#### DEFINITIONS

##### Column sum vector s

We define the vector $\mathbf{s}$ as the column sums of $\mathbf{M}$, with entries $\alpha$

$$s_\alpha := \sum_\iota M^\iota_\alpha \quad , \tag{1}$$

which by definition have non-zero entries only for the transport reactions, the ones that do not conserve internal mass.

##### Kinetic rate laws

The turnover times $\tau_\alpha$ following the Michaelis-Menten rate law [1] (as in model L3) are defined as

$$\tau_\alpha(\mathbf{a}, \rho\,\mathbf{b}) = \frac{1}{k_{\rm cat}^\alpha} \prod_\epsilon \left(1 + \frac{K_\alpha^\epsilon}{a^\epsilon}\right) \prod_\iota \left(1 + \frac{K_\alpha^\iota}{\rho\, b^\iota}\right) \quad , \tag{2}$$

where $k_{\rm cat}^\alpha$ is the corresponding turnover number (in mass of product per mass of protein catalyst per hour, resulting in $h^{-1}$), $K_\alpha^\epsilon$ (in $gL^{-1}$) are the Michaelis constants (in $gL^{-1}$) for each external reactant $\epsilon$ in reaction $\alpha$, and $K_\alpha^\iota$ are the Michaelis constants (in $gL^{-1}$) for each internal reactant $\iota$ in reaction $\alpha$. Here we use the convention that $K_\alpha^\epsilon$ and $K_\alpha^\iota$ are zero if the corresponding $\epsilon$ or $\iota$ is not a reactant in reaction $\alpha$.

The turnover times $\tau_\alpha$ following the Michaelis-Menten rate law with activation [1] (as in model G5) are defined as

$$\tau_\alpha(\mathbf{a}, \rho\,\mathbf{b}) = \frac{1}{k_{\rm cat}^\alpha} \prod_\epsilon \left(1 + \frac{K_\alpha^\epsilon}{a^\epsilon}\right) \prod_\iota \left(1 + \frac{K_\alpha^\iota}{\rho\, b^\iota}\right) \left(1 + \frac{KA_\alpha^\iota}{\rho\, b^\iota}\right) \quad , \tag{3}$$

where $KA_\alpha^\iota$ is the activation constant (in $gL^{-1}$) for the internal reactant $\iota$ in reaction $\alpha$.

#### DERIVATIONS

##### The continuity equation

Using definition (1), the biomass sum $\sum \mathbf{b} = 1$ (equation (8) in the main text) in terms of $\mathbf{q}$ as defined by $\mathbf{Mq} = \mathbf{b}$ (equation (12) in the main text) becomes

$$\sum_\iota b_\iota = 1 \quad \Rightarrow \quad \sum_{\iota,\alpha} M_\alpha^\iota\, q_\alpha = \sum_\alpha \left(\sum_\iota M_\alpha^\iota\right) q_\alpha = \sum_\alpha s_\alpha q_\alpha = 1 \quad , \tag{4}$$

which is the continuity equation (18) in the main text.

##### Ribosome fractions $\chi$

Each $\chi_\alpha$ quantifies the fraction of ribosomes allocated to produce protein $\alpha$. Thus, for each $\alpha$, mass conservation dictates that for a total protein production flux $M_{\rm r}^{\rm p} v_{\rm r}$ and protein $\alpha$ concentration $\rho\, b_{\rm p}\phi_\alpha$

$$\underbrace{\chi_\alpha M_{\rm r}^{\rm p} v_{\rm r}}_{\substack{\text{protein } \alpha \\ \text{production flux}}} = \underbrace{\mu\rho\, b_{\rm p}\phi_\alpha}_{\substack{\text{dilution} \\ \text{by growth}}} + \underbrace{\rho\, \dot{b}_{\rm p}\phi_\alpha + \rho\, b_{\rm p}\dot{\phi}_\alpha}_{\substack{\text{concentration} \\ \text{change}}} \quad . \tag{5}$$

Solving for $\chi_\alpha$

$$\chi_\alpha = \frac{\mu\rho\, b_{\rm p}\phi_\alpha + \rho\, \dot{b}_{\rm p}\phi_\alpha + \rho\, b_{\rm p}\dot{\phi}_\alpha}{M_{\rm r}^{\rm p}\, v_{\rm r}} \quad . \tag{6}$$

Substituting $b_{\rm p}$ and $\dot{b}_{\rm p}$ with equations (12,13) of the main text, respectively

$$\chi_\alpha = \frac{\mu\rho\, M_{\rm r}^{\rm p} q_{\rm r}\, \phi_\alpha + \rho\, M_{\rm r}^{\rm p} \dot{q}_{\rm r}\, \phi_\alpha + \rho\, M_{\rm r}^{\rm p} q_{\rm r}\, \dot{\phi}_\alpha}{\rho M_{\rm r}^{\rm p}\, (\mu\, q_{\rm r} + \dot{q}_{\rm r})} \quad . \tag{7}$$

Simplifying equation (7), we get

$$\chi_\alpha = \frac{\mu q_{\rm r}\, \phi_\alpha + \dot{q}_{\rm r}\, \phi_\alpha + q_{\rm r}\, \dot{\phi}_\alpha}{\mu\, q_{\rm r} + \dot{q}_{\rm r}} = \phi_\alpha + \frac{q_{\rm r}\, \dot{\phi}_\alpha}{\mu\, q_{\rm r} + \dot{q}_{\rm r}} \quad . \tag{8}$$

**The equations of optimal motion (EOM)**

Let's remind the Lagrangian $\mathcal{L}$ defined on the equation (19) in the main text

$$\mathcal{L}(t,\mathbf{q},\dot{\mathbf{q}}) := \mu(\mathbf{a}(t),\mathbf{q},\dot{\mathbf{q}}) + \lambda(\mathbf{s}\cdot\mathbf{q}-1) \quad .$$

We have the corresponding the Euler-Lagrange equations for each reaction $\alpha$

$$\frac{\partial \mathcal{L}}{\partial q^\alpha}\Big|_{\dot{\mathbf{q}}} - \frac{\mathrm{d}}{\mathrm{d}t}\frac{\partial \mathcal{L}}{\partial \dot{q}^\alpha}\Big|_{\mathbf{q}} = 0 \quad , \tag{9}$$

where, for clarity, our notation highlights that the first partial derivative is calculated at fixed $\dot{\mathbf{q}}$, and de second partial derivative is calculated at fixed $\mathbf{q}$. Let's first note that

$$\frac{\partial \mathcal{L}}{\partial q^\alpha} = \frac{\partial \mu}{\partial q^\alpha} + \lambda\, s_\alpha \tag{10}$$

and

$$\frac{\mathrm{d}}{\mathrm{d}t}\frac{\partial \mathcal{L}}{\partial \dot{q}^\alpha} = \frac{\mathrm{d}}{\mathrm{d}t}\frac{\partial \mu}{\partial \dot{q}^\alpha} \quad , \tag{11}$$

so the Euler-Lagrange equation for each reaction $\alpha$ can also be written as

$$\frac{\partial \mu}{\partial q^\alpha} + \lambda\, s_\alpha - \frac{\mathrm{d}}{\mathrm{d}t}\frac{\partial \mu}{\partial \dot{q}^\alpha} = 0 \quad . \tag{12}$$

To account now for the continuity constraint and find the optimal $\lambda$, we first multiply all terms by $q^\alpha$ (the upper index indicates there is also now a summation on the index $\alpha$)

$$\frac{\partial \mu}{\partial q^\alpha} q^\alpha + \lambda\, s_\alpha\, q^\alpha - \frac{\mathrm{d}}{\mathrm{d}t}\frac{\partial \mu}{\partial \dot{q}^\alpha} q^\alpha = 0 \quad . \tag{13}$$

We may now solve for the optimal $\lambda$ considering the continuity constraint $s_\alpha\, q^\alpha = 1$

$$\lambda = -\left(\frac{\partial \mu}{\partial q^\alpha} - \frac{\mathrm{d}}{\mathrm{d}t}\frac{\partial \mu}{\partial \dot{q}^\alpha}\right) q^\alpha \quad . \tag{14}$$

To keep the notation in the next steps more clear, it is now convenient to define the Jacobian matrix $\mathbf{J}(\mathbf{q}) := \partial\boldsymbol{\tau}/\partial\mathbf{q}$ of the turnover times $\boldsymbol{\tau}$, calculated via the chain rule

$$J_\alpha^\beta := \frac{\partial \tau^\beta}{\partial q^\alpha} = \sum_\iota \frac{\partial \tau^\beta}{\partial b^\iota}\frac{\partial b^\iota}{\partial q^\alpha} = \sum_\iota \frac{\partial \tau^\beta}{\partial b^\iota} M_\alpha^\iota \quad , \tag{15}$$

where $\partial\tau^\beta/\partial b^\iota$ is understood to be a function of $\mathbf{q}$ by substituting its dependence on $\mathbf{b}$ with the equation $\mathbf{b} = \mathbf{M}\mathbf{q}$ (equation 12 in the main text). We now calculate the following derivatives

$$\frac{\partial \mu}{\partial q^\alpha} = \frac{M_\alpha^{\mathrm{p}} - \dot{q}_\beta J_\alpha^\beta}{\mathbf{q}\cdot\boldsymbol{\tau}} - \frac{\mu(\tau_\alpha + \dot{q}_\beta J_\alpha^\beta)}{\mathbf{q}\cdot\boldsymbol{\tau}} \quad , \tag{16}$$

$$\frac{\partial \mu}{\partial \dot{q}^\alpha} = -\frac{\tau_\alpha}{\mathbf{q}\cdot\boldsymbol{\tau}} \tag{17}$$

$$\frac{\mathrm{d}}{\mathrm{d}t}\frac{\partial \mu}{\partial \dot{q}^\alpha} = -\frac{\dot{\tau}_\alpha}{\mathbf{q}\cdot\boldsymbol{\tau}} + \frac{\tau_\alpha\,(\dot{\mathbf{q}}\cdot\boldsymbol{\tau} + \mathbf{q}\cdot\dot{\boldsymbol{\tau}})}{(\mathbf{q}\cdot\boldsymbol{\tau})^2} \tag{18}$$

Substituting equations (16,18) into the equation for the optimal $\lambda$ (14)

$$\lambda = -\left(\frac{M_\alpha^{\mathrm{p}} - \dot{q}_\beta J_\alpha^\beta - \mu(\tau_\alpha + q_\beta J_\alpha^\beta) + \dot{\tau}_\alpha}{\mathbf{q}\cdot\boldsymbol{\tau}} - \frac{\tau_\alpha\,(\dot{\mathbf{q}}\cdot\boldsymbol{\tau} + \mathbf{q}\cdot\dot{\boldsymbol{\tau}})}{(\mathbf{q}\cdot\boldsymbol{\tau})^2}\right) q^\alpha \quad . \tag{19}$$

Multiplying by $q^\alpha$ (with implied summation on the index $\alpha$)

$$\lambda = -\left(\frac{M^{\rm p}_{\rm r}\, q_{\rm r} - \dot{\mathbf{q}}^\top \mathbf{J}\,\mathbf{q} - \mu(\mathbf{q}\cdot\boldsymbol{\tau} + \mathbf{q}^\top \mathbf{J}\,\mathbf{q}) + \mathbf{q}\cdot\dot{\boldsymbol{\tau}} - \mathbf{q}\cdot\dot{\boldsymbol{\tau}} - \dot{\mathbf{q}}\cdot\boldsymbol{\tau}}{\mathbf{q}\cdot\boldsymbol{\tau}}\right) \quad , \tag{20}$$

where we simplified $M^{\rm p}_\alpha\, q^\alpha = M^{\rm p}_{\rm r}\, q_{\rm r}$ since $M^{\rm p}_{\rm r}$ is the only non-zero entry in the row $M^{\rm p}_\alpha$ by definition. Canceling the term $\mathbf{q}\cdot\dot{\boldsymbol{\tau}}$

$$\lambda = -\left(\frac{M^{\rm p}_{\rm r}\, q_{\rm r} - \dot{\mathbf{q}}^\top \mathbf{J}\,\mathbf{q} - \mu(\mathbf{q}\cdot\boldsymbol{\tau} + \mathbf{q}^\top \mathbf{J}\,\mathbf{q}) - \dot{\mathbf{q}}\cdot\boldsymbol{\tau}}{\mathbf{q}\cdot\boldsymbol{\tau}}\right) \quad . \tag{21}$$

Rearranging,

$$\lambda = -\left(\frac{M^{\rm p}_{\rm r}\, q_{\rm r} - \dot{\mathbf{q}}\cdot\boldsymbol{\tau} - \mu\,\mathbf{q}\cdot\boldsymbol{\tau}}{\mathbf{q}\cdot\boldsymbol{\tau}} - \frac{\dot{\mathbf{q}}^\top \mathbf{J}\,\mathbf{q} + \mu\,\mathbf{q}^\top \mathbf{J}\,\mathbf{q}}{\mathbf{q}\cdot\boldsymbol{\tau}}\right) \quad . \tag{22}$$

We now note the first term in parentheses is zero due to the definition of growth rate (equation (17) in the main text), so the *optimal* $\lambda$

$$\lambda = \frac{\left(\dot{\mathbf{q}}^\top + \mu\,\mathbf{q}^\top\right)\mathbf{J}\,\mathbf{q}}{\mathbf{q}\cdot\boldsymbol{\tau}} \quad . \tag{23}$$

Now substituting equations (16,18,23) into the Euler-Lagrange equation (12)

$$\frac{M^{\rm p}_\alpha - \dot{q}_\beta J^\beta_\alpha}{\mathbf{q}\cdot\boldsymbol{\tau}} - \frac{\mu(\tau_\alpha + q_\beta J^\beta_\alpha)}{\mathbf{q}\cdot\boldsymbol{\tau}} + \frac{s_\alpha\left(\dot{\mathbf{q}}^\top + \mu\,\mathbf{q}^\top\right)\mathbf{J}\,\mathbf{q}}{\mathbf{q}\cdot\boldsymbol{\tau}} + \frac{\dot{\tau}_\alpha}{\mathbf{q}\cdot\boldsymbol{\tau}} - \frac{\tau_\alpha\left(\dot{\mathbf{q}}\cdot\boldsymbol{\tau} + \mathbf{q}\cdot\dot{\boldsymbol{\tau}}\right)}{\left(\mathbf{q}\cdot\boldsymbol{\tau}\right)^2} = 0 \tag{24}$$

Multiplying by $\mathbf{q}\cdot\boldsymbol{\tau}$

$$M^{\rm p}_\alpha - \dot{q}_\beta J^\beta_\alpha - \mu(\tau_\alpha + q_\beta J^\beta_\alpha) + s_\alpha\left(\dot{\mathbf{q}}^\top + \mu\,\mathbf{q}^\top\right)\mathbf{J}\,\mathbf{q} + \dot{\tau}_\alpha - \frac{\tau_\alpha\left(\dot{\mathbf{q}}\cdot\boldsymbol{\tau} + \mathbf{q}\cdot\dot{\boldsymbol{\tau}}\right)}{\mathbf{q}\cdot\boldsymbol{\tau}} = 0 \tag{25}$$

Rearranging, we get equation (21) in the main text. We calculate the time derivatives $\dot{\tau}_\alpha$ via the chain rule

$$\dot{\tau}_\alpha = \frac{\partial \tau_\alpha}{\partial q^\beta}\frac{{\rm d}q^\beta}{{\rm d}t} + \frac{\partial \tau_\alpha}{\partial a^\epsilon}\frac{{\rm d}a^\epsilon}{{\rm d}t} = \frac{\partial \tau_\alpha}{\partial b^\iota} M^\iota_\beta\, \dot{q}^\beta + \frac{\partial \tau_\alpha}{\partial a^\epsilon}\dot{a}^\epsilon \tag{26}$$

#### SI FIGURES

##### Figures relating to GCMs L3 and G5 in the main text

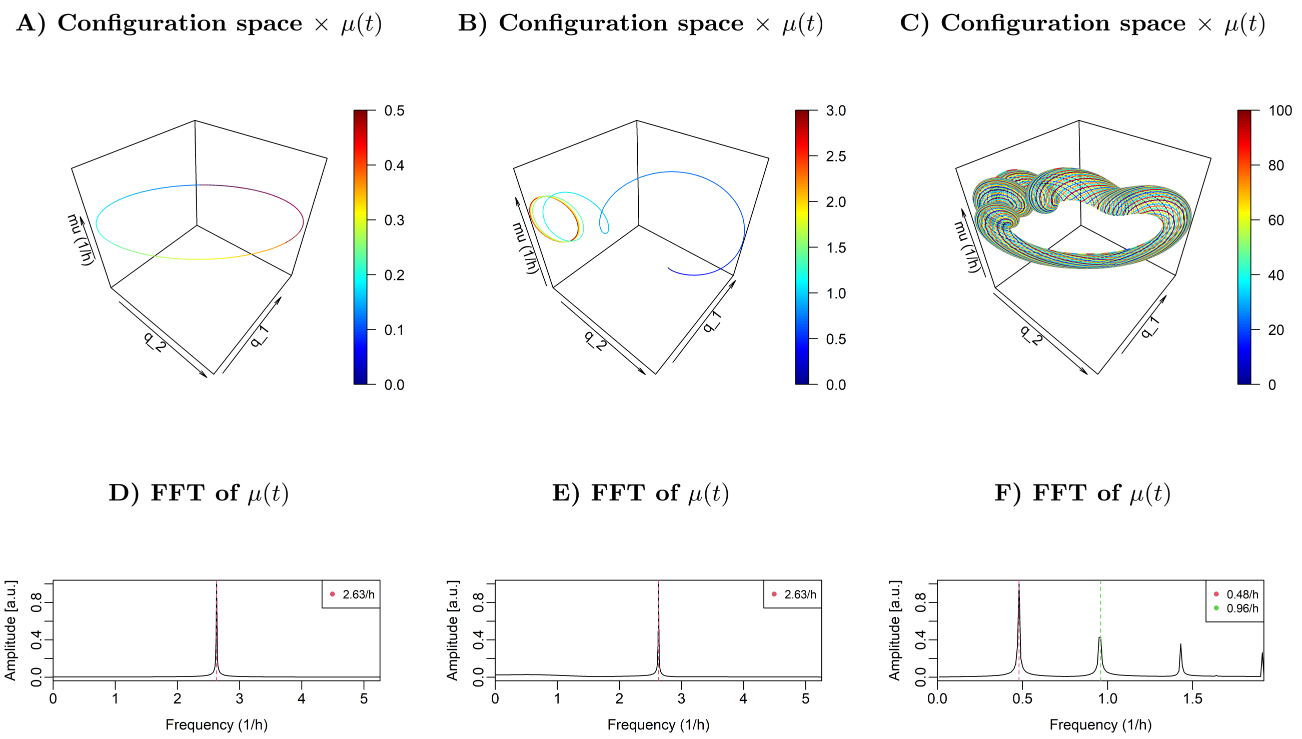


FIG. S1: The OGTs in the configuration space $\times$ $\mu(t)$ (A,B,C) for model L3 at the corresponding three different media and initial conditions of Fig.2 in the main text, and the respective FFT of instantaneous growth rate $\mu(t)$ (D,E,F). For A, B, C, the color bar indicates the simulation time in hours. In (A) we observe a closed orbit, which is consistent with the single fundamental frequency ($\nu = 2.63\ h^{-1}$) indicated in the corresponding FFT in (D). In (B) we see the system goes through a transition phase and then also converges to a closed orbit (red color) with single frequency ($\nu = 2.63\ h^{-1}$) indicated on (E). On (C) the trajectory indicates a more complicated dynamics resembling a limit torus, which is consistent with the two incommensurable fundamental frequencies of the medium oscillation ($\nu_{\mathrm{m}} \approx 0.48\ h^{-1}$) and natural frequency of the model $\nu \approx 2.6\ h^{-1}$ observed in its corresponding FFT of $\mu(t)$ in (F), together with some of their harmonics.

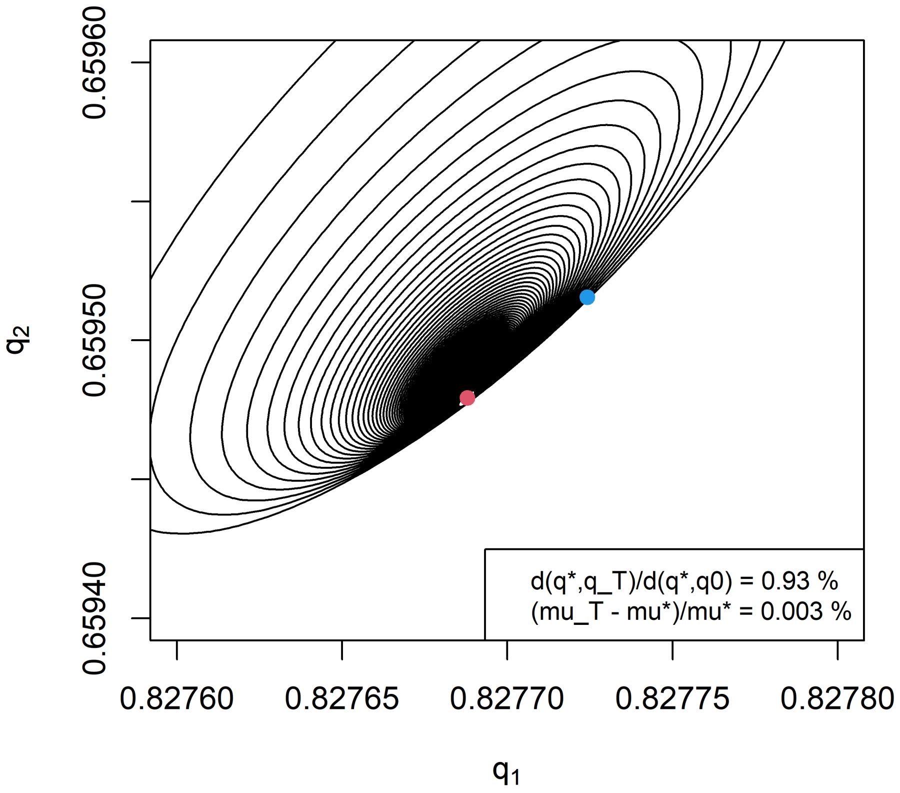


FIG. S2: The time average state in time $\bar{q}(t)$ (black line) of model L3 at static medium $a_C = 10\ g\ L^{-1}$ converges to a point (red dot) very close (more than 100 times closer than the initial $\mathbf{q}_0$), but different than the OGS (blue dot) of this model in the same medium. Here the black lines are the final 98h of a 100h simulation. The corresponding average growth rate $\bar{\mu}(t)$ also converges to a value slightly higher (0.003 %) than the $\mu^*$ at the OGS.

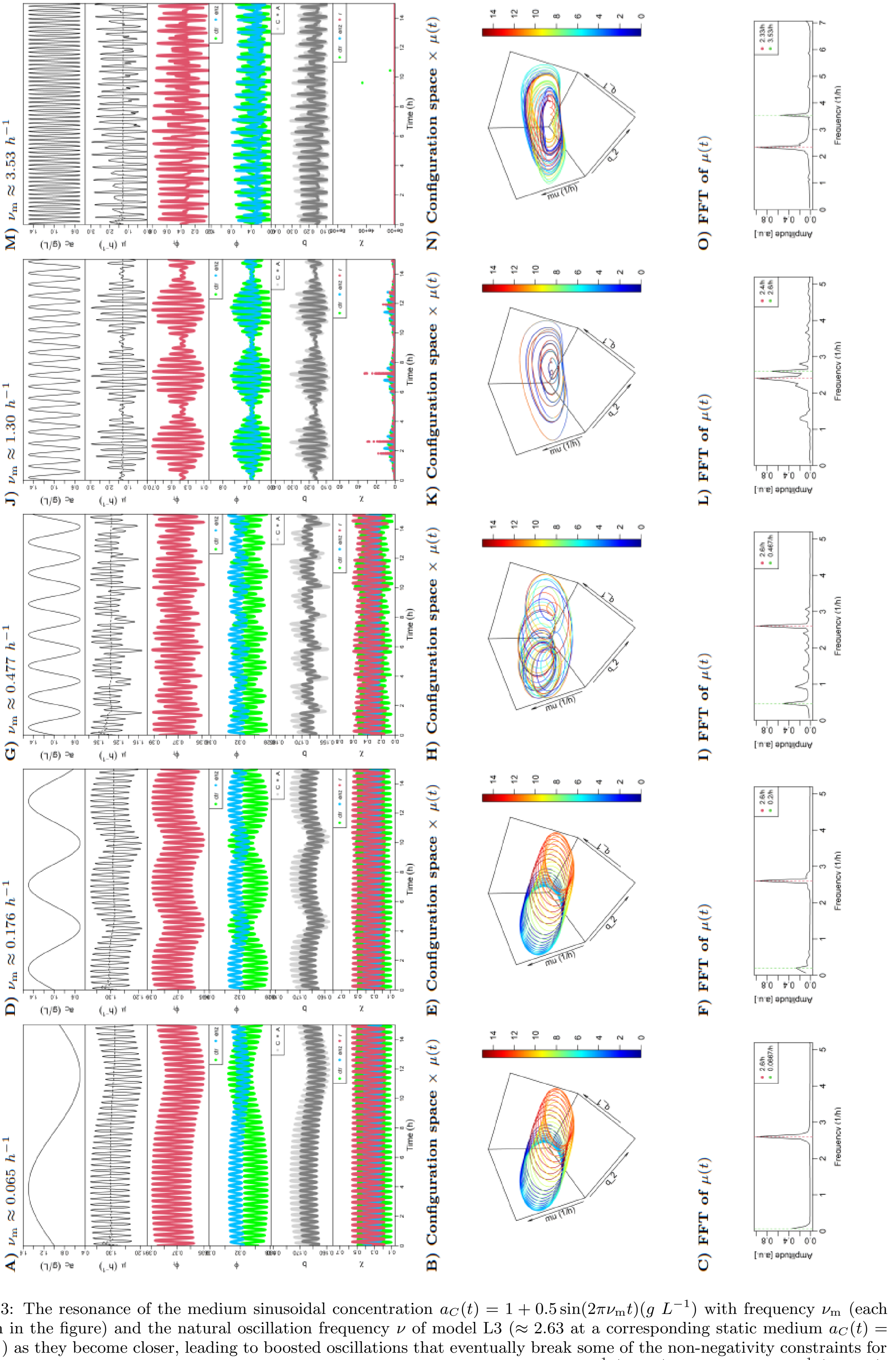


FIG. S3: The resonance of the medium sinusoidal concentration $a_C(t) = 1 + 0.5\sin(2\pi\nu_\mathrm{m}t)(g\ L^{-1})$ with frequency $\nu_\mathrm{m}$ (each column in the figure) and the natural oscillation frequency $\nu$ of model L3 ($\approx 2.63$ at a corresponding static medium $a_C(t) = 1g\ L^{-1}$) as they become closer, leading to boosted oscillations that eventually break some of the non-negativity constraints for proteome allocation $\boldsymbol{\phi}$, biomass allocation $\mathbf{b}$ and ribosome allocation $\boldsymbol{\chi}$ at $\nu_\mathrm{m} \approx 1.30\ h^{-1}$ (J,K,L) and $\nu_\mathrm{m} \approx 3.53\ h^{-1}$ (M,N,O).

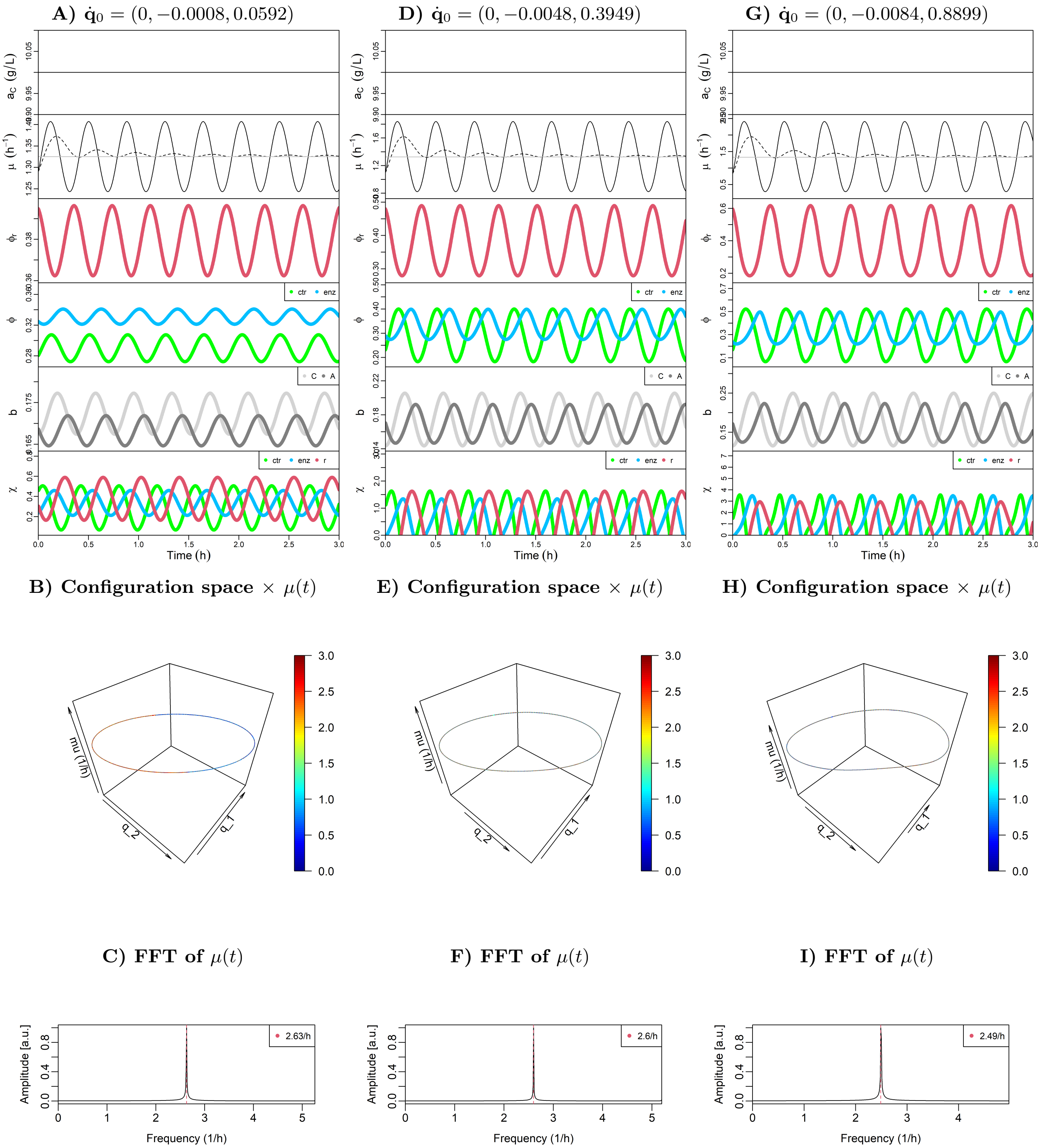


FIG. S4: The effect of extreme initial velocities $\dot{\mathbf{q}}_0$ (different values for each column) for model L3 in the same static medium $a_C = 10\ g\ L^{-1}$, as shown by the cell state properties (A,D,G), their trajectories in the configuration space $\times\ \mu(t)$ (B,E,H) and the corresponding FFT of $\mu(t)$ (C,F,I). In (A,B,C) we see the limit $\dot{\mathbf{q}}_0$ where the ribosome fractions $\boldsymbol{\chi}$ remain non-negative, while in the other cases not. For the extreme initial velocity $\dot{\mathbf{q}}_0$ in (G,H,I) we see that oscillations in the cell state variables (G) do not seem anymore as simple sine waves, and the trajectory in (H) more clearly deviates from a simple elliptic orbit.

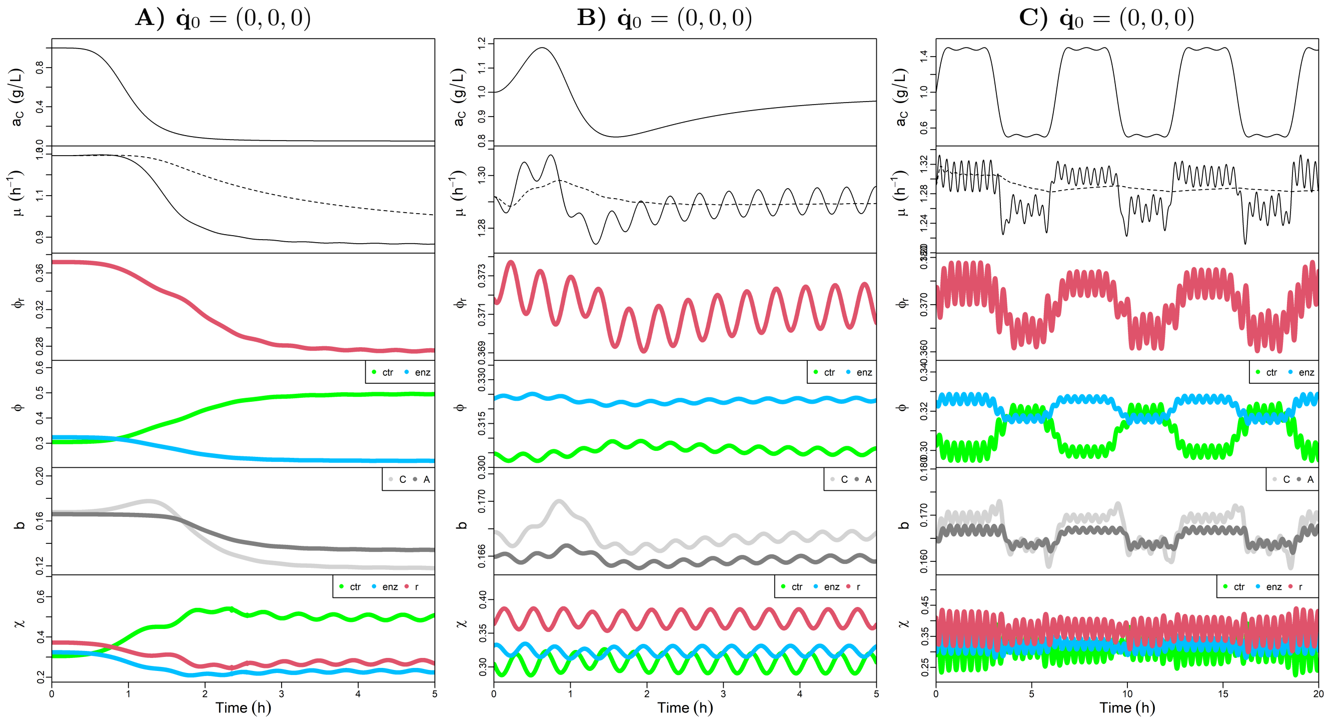


FIG. S5: Simulations of model L3 on three different media shifts. **A)** Medium downshift defined by the function $a_C(t) = 1 - 0.95\frac{t^5}{t^5+1}$ $(g\ L^{-1})$. **B)** Medium upshift followed by downshift defined by the function $a_C(t) = 1 - 0.95\frac{t^5}{t^5+1} + 0.95\frac{t^2}{t^2+1}$ $(g\ L^{-1})$. **C)** Periodic medium emulating feast-and-famine shifts with the function $a_C(t) = 1 + 0.5sin(t + sin(2t))$ $(g\ L^{-1})$, see next Fig. S6A for a corresponding longer simulation with $T = 100\ h$.

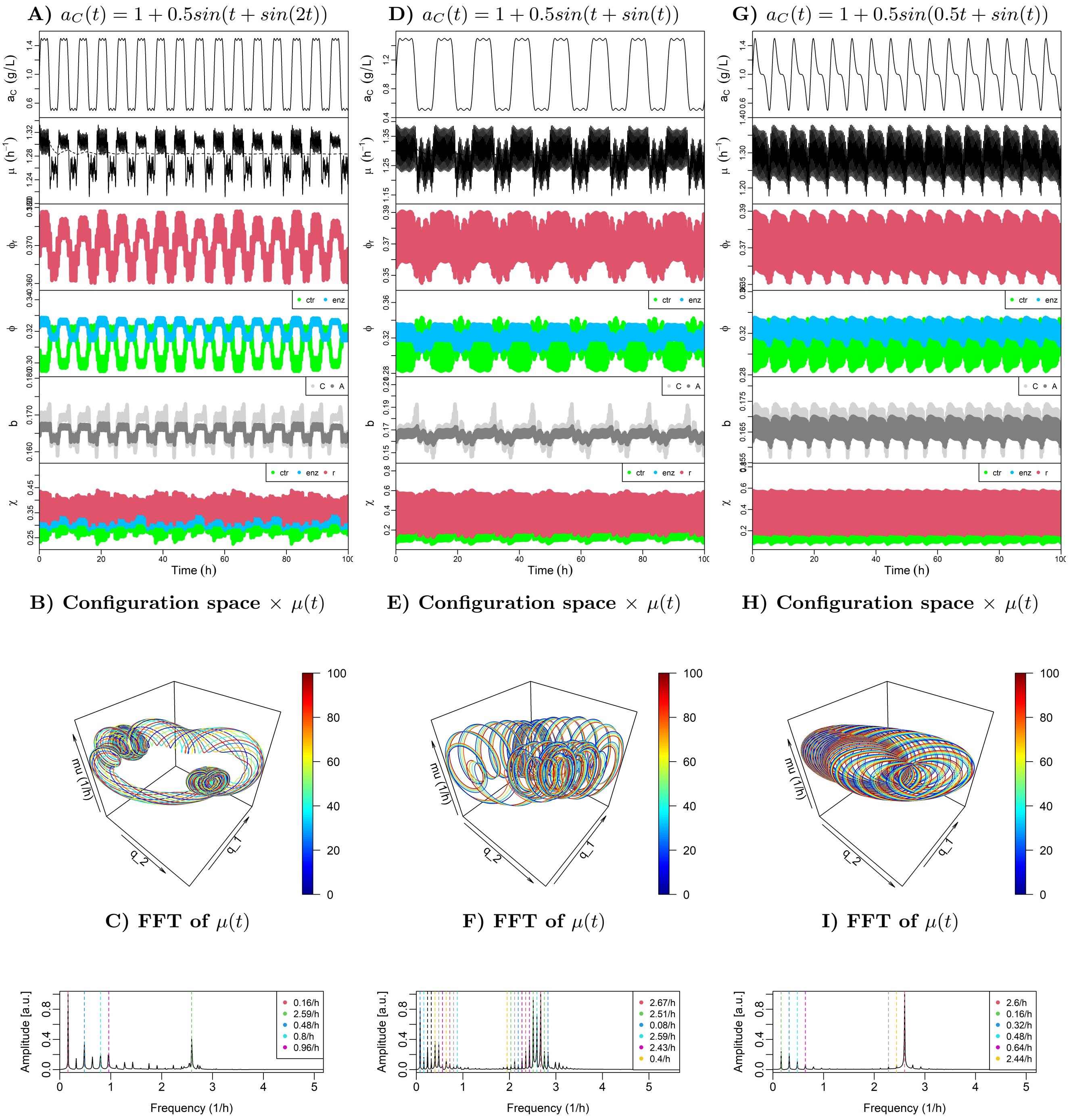


FIG. S6: Longer simulations ($T = 100$ $h$) for the model L3 on different periodic media (in each column).

$$\mathbf{K} = \begin{array}{c} \begin{matrix} [\text{ctr}] & [\text{met}] & [\text{DNAp}] & [\text{RNAp}] & [\text{r}] \end{matrix} \\ \left[ \begin{matrix} 0.1 & 0 & 0 & 0 & 0 \\ 0 & 11 & 0 & 0 & 0 \\ 0 & 0 & 0 & 0 & 15 \\ 0 & 0 & 10 & 10 & 0 \\ 0 & 0 & 0 & 0 & 29 \end{matrix} \right] \end{array} \begin{matrix} \{C_{ex}\} \\ (C) \\ (ATP) \\ (NT) \\ (AA) \end{matrix}$$

$$\mathbf{KA} = \begin{array}{c} \begin{matrix} [\text{ctr}] & [\text{met}] & [\text{DNAp}] & [\text{RNAp}] & [\text{r}] \end{matrix} \\ \left[ \begin{matrix} 0 & 0 & 5 & 5 & 0 \\ 0 & 0 & 0 & 0 & 35 \end{matrix} \right] \end{array} \begin{matrix} (DNA) \\ (RNA) \end{matrix}$$

$$\mathbf{k}_{\text{cat}} = \begin{array}{c} \begin{matrix} [\text{ctr}] & [\text{met}] & [\text{DNAp}] & [\text{RNAp}] & [\text{r}] \end{matrix} \\ \left[ \begin{matrix} 12 & 17 & 5 & 5 & 12 \end{matrix} \right] \end{array}$$

$$\rho = 340$$

FIG. S7: Parameters defining Model G5, in addition to the matrix **M** on Fig.(3) in the main text and the rate law with activation (3). Note here DNA and RNA are produced but not consumed by any reaction; instead, DNA production is necessary to activate it own production and the production of RNA, and RNA is necessary to activate the ribosome reaction producing proteins.

**A) Configuration space $\times$ $\mu(t)$**

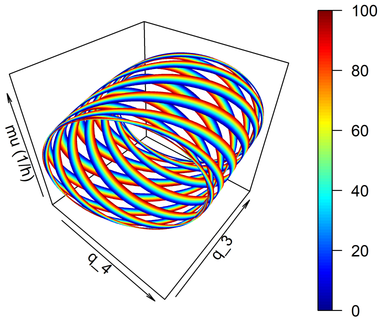


**B) FFT of $\mu(t)$**

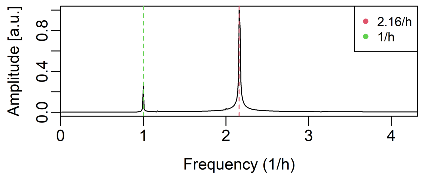


FIG. S8: The OGT in the configuration space $\times$ $\mu(t)$ (A) for model G5 at the corresponding medium and initial conditions of Fig.3 in the main text, and the respective FFT of instantaneous growth rate $\mu(t)$ (B).

**More GCM examples: L2, L4, Y4, L5, G9.**

$$\mathbf{M} = \begin{array}{c} \\ \\ \end{array}\begin{array}{cc} [\text{ctr}] & [\text{r}] \\ \left[\begin{array}{c} 1 \\ 0 \end{array}\right. & \left.\begin{array}{c} -1 \\ 1 \end{array}\right] \end{array}\begin{array}{c} \\ (C) \\ (p) \end{array}$$

$$\mathbf{K} = \begin{array}{cc} [\text{ctr}] & [\text{r}] \\ \left[\begin{array}{c} 0.1 \\ 0 \\ 0 \end{array}\right. & \left.\begin{array}{c} 0 \\ 40 \\ 0 \end{array}\right] \end{array}\begin{array}{c} \\ \{C_{\text{ex}}\} \\ (C) \\ (p) \end{array}$$

$$\mathbf{k}_{\text{cat}} = \begin{array}{cc} [\text{ctr}] & [\text{r}] \\ [\ 6 & 4\ ] \end{array}$$

$$\rho = 340$$

FIG. S9: Model L2: a CGM with the simplest possible structure including only two reactions following the Michaelis-Menten kinetics (2).

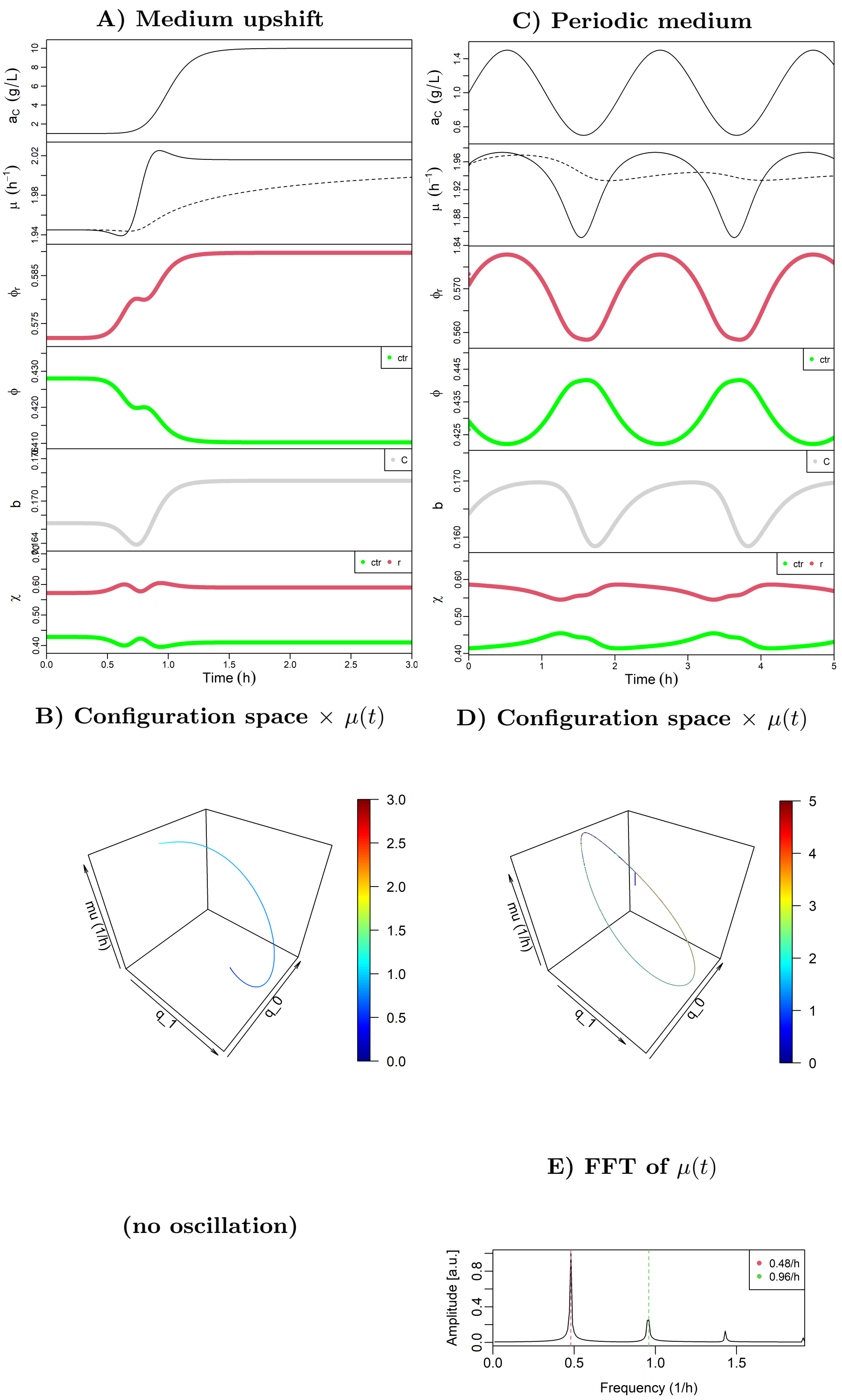


FIG. S10: Numerical solutions for the equations of optimal motion of model L2 described in Fig.(S9), on a medium upshift (left column) and periodic medium (right column).

$$\mathbf{M} = \begin{array}{c} \begin{matrix} [\text{ctr}] & [\text{enz1}] & [\text{enz2}] & [\text{r}] \end{matrix} \\ \left[\begin{matrix} 1 & 0 & 0 & 0 \\ 0 & -1 & 0 & 0 \\ 0 & 1 & -1 & -1 \\ 0 & 0 & 0 & 1 \end{matrix}\right] \end{array} \begin{matrix} (C) \\ (C2) \\ (A) \\ (p) \end{matrix}$$

$$\mathbf{K} = \begin{array}{c} \begin{matrix} [\text{ctr}] & [\text{enz1}] & [\text{enz2}] & [\text{r}] \end{matrix} \\ \left[\begin{matrix} 0.1 & 0 & 0 & 0 \\ 0 & 14 & 0 & 0 \\ 0 & 0 & 26 & 0 \\ 0 & 0 & 0 & 35 \\ 0 & 0 & 0 & 0 \end{matrix}\right] \end{array} \begin{matrix} C_{ex}\} \\ (C) \\ (N) \\ (AA) \\ (p) \end{matrix}$$

$$\mathbf{k}_{\text{cat}} = \begin{array}{c} \begin{matrix} [\text{ctr}] & [\text{enz1}] & [\text{enz2}] & [\text{r}] \end{matrix} \\ \left[\begin{matrix} 9 & 17 & 13 & 6 \end{matrix}\right] \end{array}$$

$$\rho = 340$$

FIG. S11: Model L4, defined with four Michaelis-Menten reactions (2) in a linear pathway.

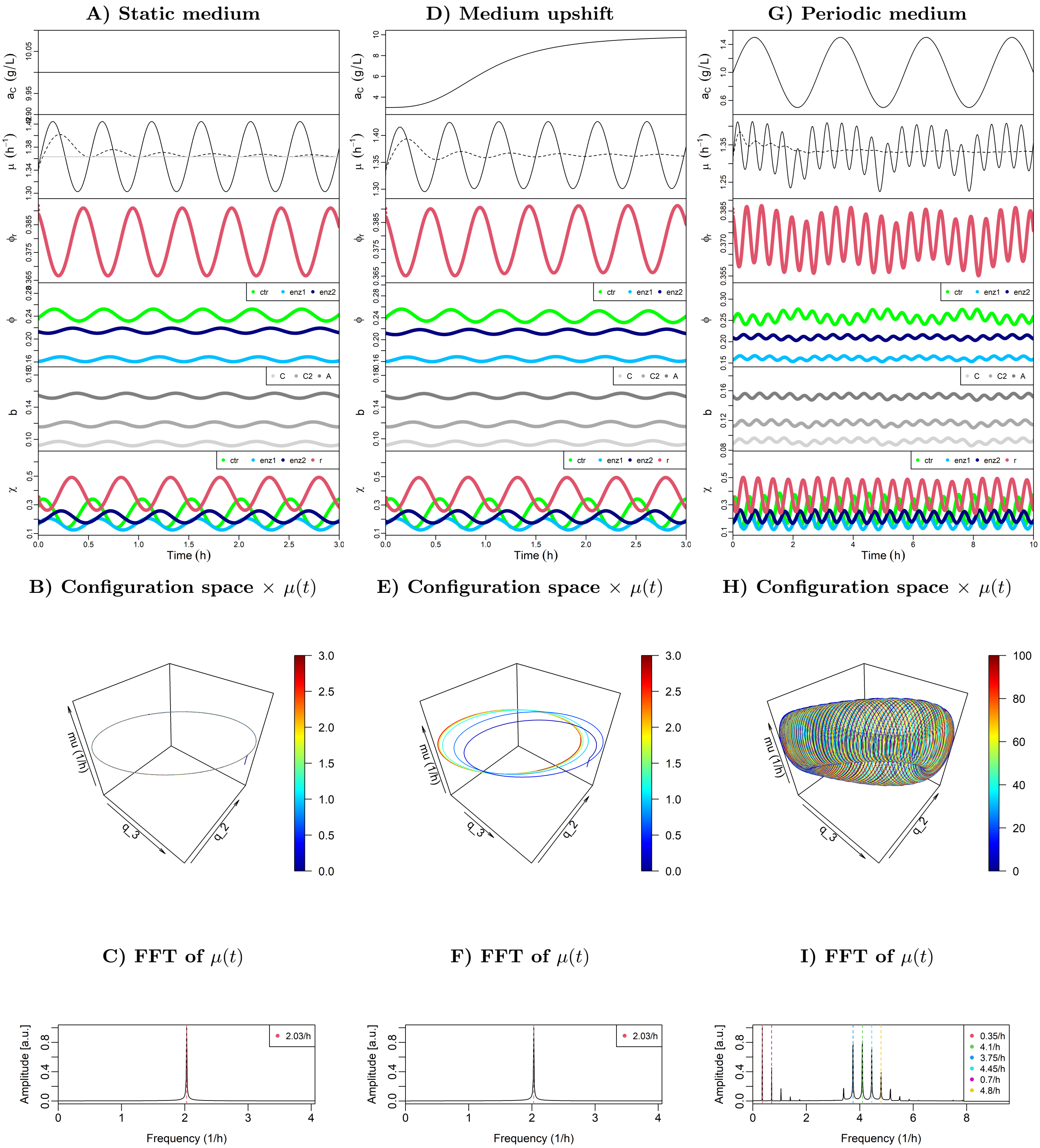


FIG. S12: Numerical solutions for the OGT of model L4 in three different media and initial conditions, as shown by the cell state properties (A,D,G), their trajectories in the configuration space $\times\ \mu(t)$ (B,E,H) and the corresponding FFT of $\mu(t)$ (C,F,I). We see the same type of qualitative behavior observed for model L3 (Fig.S1 and Fig.2 in the main text) in the same media.

$$\mathbf{M} = \begin{array}{c} \\ \left[\begin{array}{cccc} 1 & 0 & -0.8 & 0 \\ 0 & 1 & -0.2 & -0.2 \\ 0 & 0 & 1 & -0.8 \\ 0 & 0 & 0 & 1 \end{array}\right] \end{array} \begin{array}{l} \\ (C) \\ (N) \\ (AA) \\ (p) \end{array}$$

Columns: [ctr] [ntr] [met] [r]

$$\mathbf{K} = \left[\begin{array}{cccc} 0.1 & 0 & 0 & 0 \\ 0 & 0.1 & 0 & 0 \\ 0 & 0 & 22 & 0 \\ 0 & 0 & 9 & 9 \\ 0 & 0 & 0 & 39 \\ 0 & 0 & 0 & 0 \end{array}\right] \begin{array}{l} \{C_{ex}\} \\ \{N_{ex}\} \\ (C) \\ (N) \\ (AA) \\ (p) \end{array}$$

Columns: [ctr] [ntr] [met] [r]

$$\mathbf{k}_{\text{cat}} = \begin{array}{cccc} [\text{ctr}] & [\text{ntr}] & [\text{met}] & [\text{r}] \\ \hline \end{array}\left[\begin{array}{cccc} 10 & 8 & 10 & 5 \end{array}\right]$$

$$\rho = 340$$

FIG. S13: Model Y4, with four Michaelis-Menten (2) reactions in a branched network including two transporters.

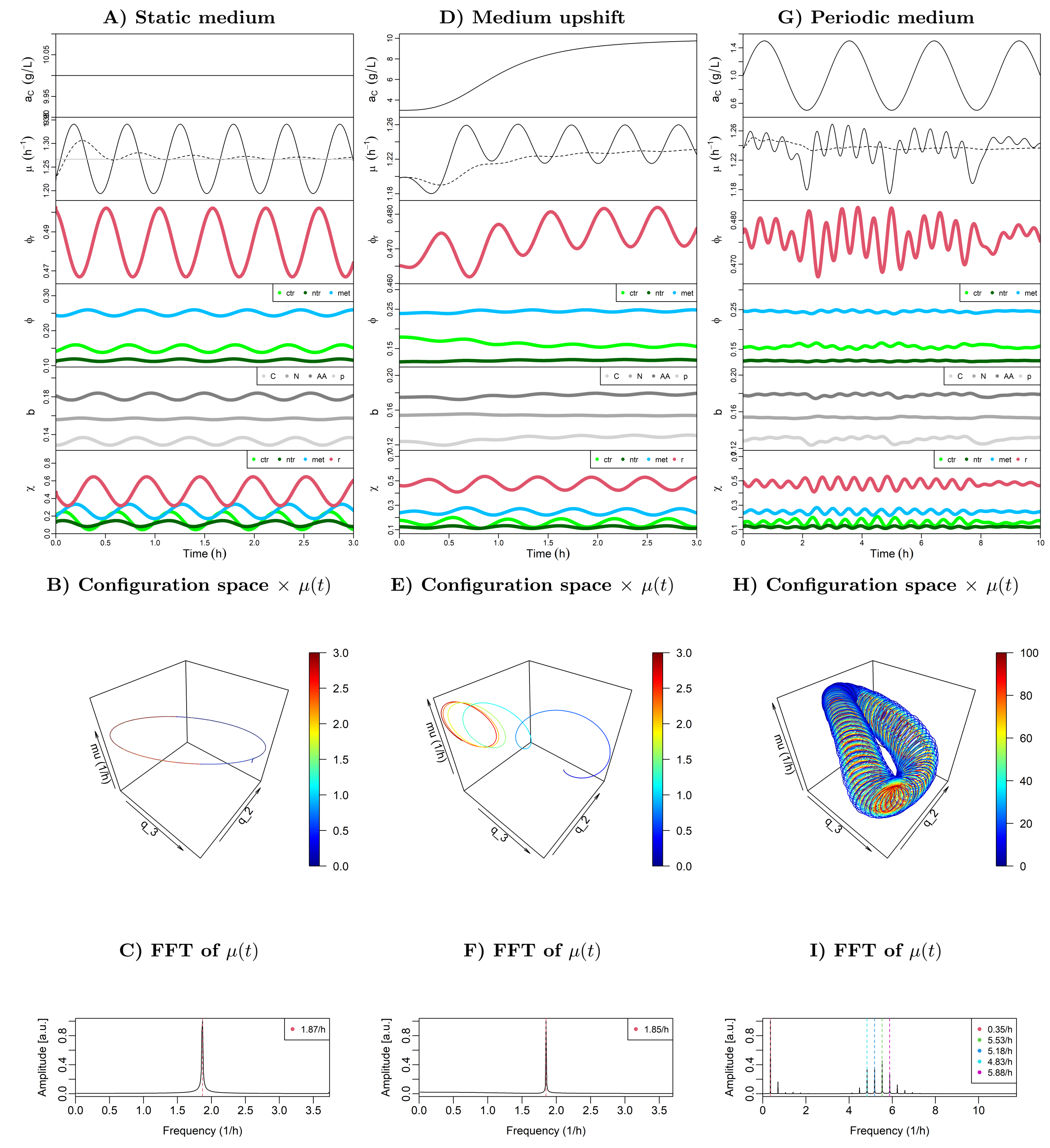


FIG. S14: Numerical solutions for the OGT of model Y4 in three different media and initial conditions, as shown by the cell state properties (A,D,G), their trajectories in the configuration space $\times\ \mu(t)$ (B,E,H) and the corresponding FFT of $\mu(t)$ (C,F,I). We see the same type of qualitative behavior observed for model L3 (Fig.S1 and Fig.2 in the main text) in the same media.

$$\mathbf{M} = \begin{array}{c} \\ \end{array}\begin{matrix} [\text{ctr}] & [\text{enz1}] & [\text{enz2}] & [\text{enz3}] & [\text{r}] & \\ \end{matrix}$$

$$\mathbf{M} = \begin{bmatrix} 1 & -1 & 0 & 0 & 0 \\ 0 & 1 & -1 & 0 & 0 \\ 0 & 0 & 1 & -1 & 0 \\ 0 & 0 & 0 & 1 & -1 \\ 0 & 0 & 0 & 0 & 1 \end{bmatrix} \begin{matrix} (C) \\ (C2) \\ (C3) \\ (A) \\ (p) \end{matrix}$$

(columns: [ctr] [enz1] [enz2] [enz3] [r])

$$\mathbf{K} = \begin{bmatrix} 0.1 & 0 & 0 & 0 & 0 \\ 0 & 10 & 0 & 0 & 0 \\ 0 & 0 & 18 & 0 & 0 \\ 0 & 0 & 0 & 25 & 0 \\ 0 & 0 & 0 & 0 & 30 \\ 0 & 0 & 0 & 0 & 0 \end{bmatrix} \begin{matrix} \{C_{ex}\} \\ (C) \\ (C2) \\ (C3) \\ (A) \\ (p) \end{matrix}$$

(columns: [ctr] [enz1] [enz2] [enz3] [r])

$$\mathbf{k}_{\text{cat}} = \begin{matrix} [\text{ctr}] & [\text{enz1}] & [\text{enz2}] & [\text{enz3}] & [\text{r}] \\ \end{matrix}$$
$$\mathbf{k}_{\text{cat}} = \begin{bmatrix} 12 & 22 & 24 & 37 & 6 \end{bmatrix}$$

$$\rho = 340$$

FIG. S15: Model L5, with 5 Michaelis-Menten (2) reactions in a linear pathway.

**A) Model L5 on static medium**

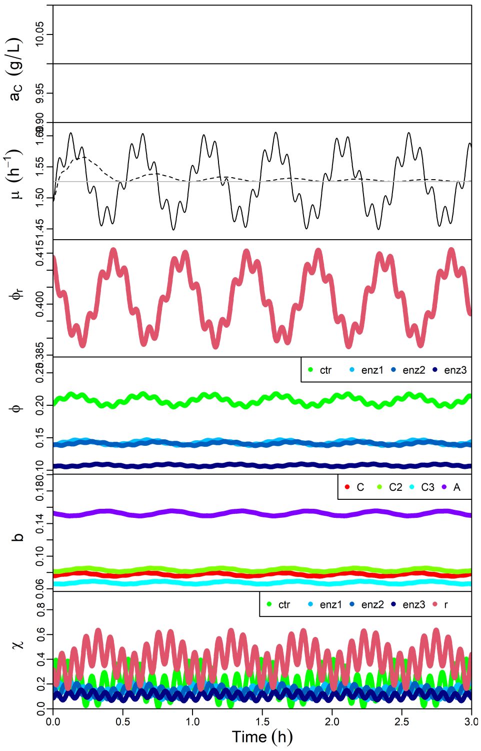


**B) Configuration space × $\mu(t)$**

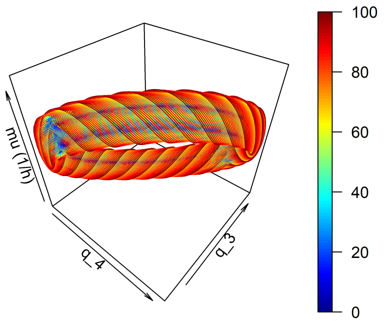


**C) FFT of $\mu(t)$**

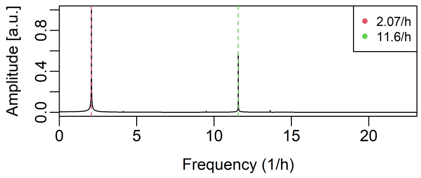


FIG. S16: Numerical solution for the equations of optimal motion of model L5 in a static medium $a_C = 10\ g\ L^{-1}$ with initial velocity $\dot{\mathbf{q}} \neq 0$, showing an apparent quasi-periodic behavior as indicated by its state variables (A), trajectory in the configuration space × $\mu(t)$ (B), and two fundamental frequencies showing in the FFT of $\mu(t)$ (C).

$\mathbf{M} =$

| | [ctr] | [ntr] | [met] | [NTS] | [AAS] | [DNAp] | [RNAp] | [TCS] | [r] |
|---|---|---|---|---|---|---|---|---|---|
| $(C)$ | 1 | 0 | −0.4 | −0.5 | −0.6 | 0 | 0 | 0 | 0 |
| $(N)$ | 0 | 1 | −0.2 | −0.3 | −0.2 | 0 | 0 | 0 | 0 |
| $(ADP)$ | 0 | 0 | −0.4 | 0.1 | 0.1 | 0 | 0 | 0.1 | 0.1 |
| $(ATP)$ | 0 | 0 | 1 | −0.2 | −0.2 | 0 | 0 | −0.1 | −0.1 |
| $(A)$ | 0 | 0 | 0 | 0 | 0.9 | 0 | 0 | −0.2 | 0 |
| $(NT)$ | 0 | 0 | 0 | 0.9 | 0 | −1 | −1 | 0 | 0 |
| $(DNA)$ | 0 | 0 | 0 | 0 | 0 | 1 | 0 | 0 | 0 |
| $(RNA)$ | 0 | 0 | 0 | 0 | 0 | 0 | 1 | −0.7 | 0.6 |
| $(TC)$ | 0 | 0 | 0 | 0 | 0 | 0 | 0 | 0.9 | −0.9 |
| $(p)$ | 0 | 0 | 0 | 0 | 0 | 0 | 0 | 0 | 0.3 |

$\mathbf{K} =$

| | [ctr] | [ntr] | [met] | [NTS] | [AAS] | [DNAp] | [RNAp] | [TCS] | [r] |
|---|---|---|---|---|---|---|---|---|---|
| $\{C_{ex}\}$ | 0.1 | 0 | 0 | 0 | 0 | 0 | 0 | 0 | 0 |
| $\{N_{ex}\}$ | 0 | 0.1 | 0 | 0 | 0 | 0 | 0 | 0 | 0 |
| $(C)$ | 0 | 0 | 4 | 4 | 4 | 0 | 0 | 0 | 0 |
| $(N)$ | 0 | 0 | 2 | 2 | 2 | 0 | 0 | 0 | 0 |
| $(ADP)$ | 0 | 0 | 17 | 0 | 0 | 0 | 0 | 0 | 0 |
| $(ATP)$ | 0 | 0 | 0 | 11 | 11 | 0 | 0 | 11 | 11 |
| $(A)$ | 0 | 0 | 0 | 0 | 0 | 0 | 0 | 6 | 0 |
| $(NT)$ | 0 | 0 | 0 | 0 | 0 | 5 | 5 | 0 | 0 |
| $(DNA)$ | 0 | 0 | 0 | 0 | 0 | 0 | 0 | 0 | 0 |
| $(RNA)$ | 0 | 0 | 0 | 0 | 0 | 0 | 0 | 7 | 0 |
| $(TC)$ | 0 | 0 | 0 | 0 | 0 | 0 | 0 | 0 | 14 |
| $(p)$ | 0 | 0 | 0 | 0 | 0 | 0 | 0 | 0 | 0 |

$\mathbf{KA} =$

| | [ctr] | [ntr] | [met] | [NTS] | [AAS] | [DNAp] | [RNAp] | [TCS] | [r] |
|---|---|---|---|---|---|---|---|---|---|
| $(C)$ | 0 | 0 | 0 | 0 | 0 | 0 | 0 | 0 | 0 |
| $(N)$ | 0 | 0 | 0 | 0 | 0 | 0 | 0 | 0 | 0 |
| $(ADP)$ | 0 | 0 | 0 | 0 | 0 | 0 | 0 | 0 | 0 |
| $(ATP)$ | 0 | 0 | 0 | 0 | 0 | 0 | 0 | 0 | 0 |
| $(A)$ | 0 | 0 | 0 | 0 | 0 | 0 | 0 | 0 | 0 |
| $(NT)$ | 0 | 0 | 0 | 0 | 0 | 0 | 0 | 0 | 0 |
| $(DNA)$ | 0 | 0 | 0 | 0 | 0 | 5 | 5 | 0 | 0 |
| $(RNA)$ | 0 | 0 | 0 | 0 | 0 | 0 | 0 | 0 | 0 |
| $(TC)$ | 0 | 0 | 0 | 0 | 0 | 0 | 0 | 0 | 0 |
| $(p)$ | 0 | 0 | 0 | 0 | 0 | 0 | 0 | 0 | 0 |

$\mathbf{k}_{\mathrm{cat}} =$

| [ctr] | [ntr] | [met] | [NTS] | [AAS] | [DNAp] | [RNAp] | [TCS] | [r] |
|---|---|---|---|---|---|---|---|---|
| 6 | 16 | 32 | 25 | 23 | 4 | 19 | 87 | 40 |

$$\rho = 340$$

FIG. S17: Model G9 parameters, assuming the rate law with activation (3).

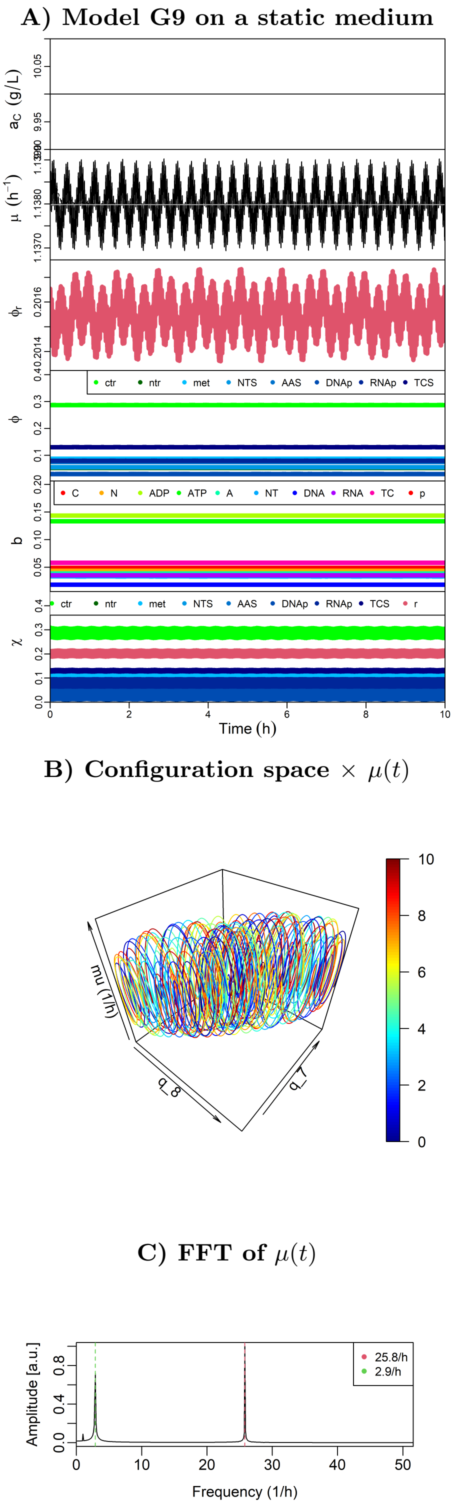


FIG. S18: Numerical solution for the equations of optimal motion of model G9 in a static medium $a_C = 10\ g\ L^{-1}$ with initial velocity $\dot{\mathbf{q}} \neq 0$, showing an apparent quasi-periodic behavior as indicated by its state variables (A), trajectory in the configuration space $\times\ \mu(t)$ (B), and three fundamental frequencies showing in the FFT of $\mu(t)$ (C).

#### Peak allocation time

##### A) Model L3 peak allocation time in a static medium

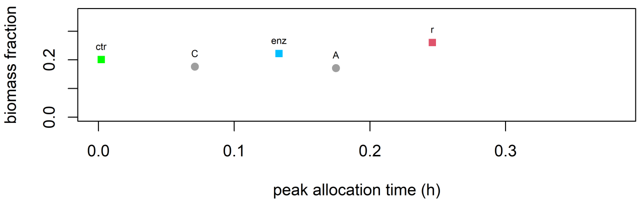


##### B) Model L4 peak allocation time in a static medium

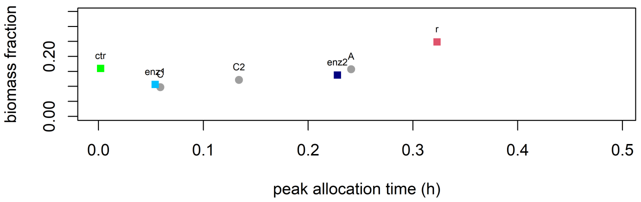


##### C) Model Y4 peak allocation time in a static medium

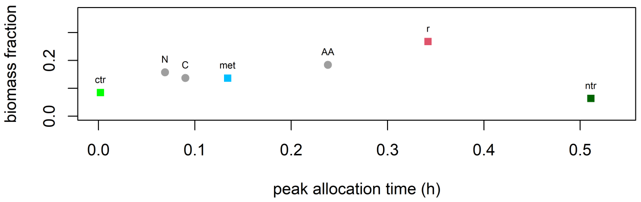


##### D) Model L5 peak allocation time in a static medium

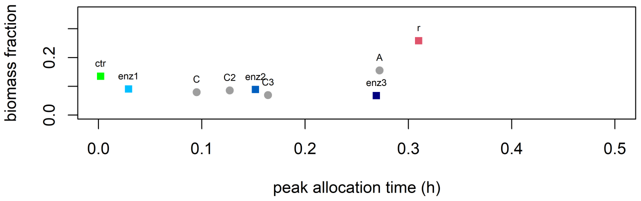


##### E) Model G9 peak allocation time in a static medium

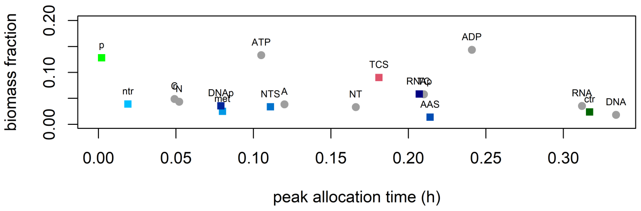


FIG. S19: Peak allocation time for models in a static medium $a_C = 10\ g\ L^{-1}$ (for models Y4 and G9 also $a_N = 10\ g\ L^{-1}$).

**Growth laws for frequency and ribosome allocation**

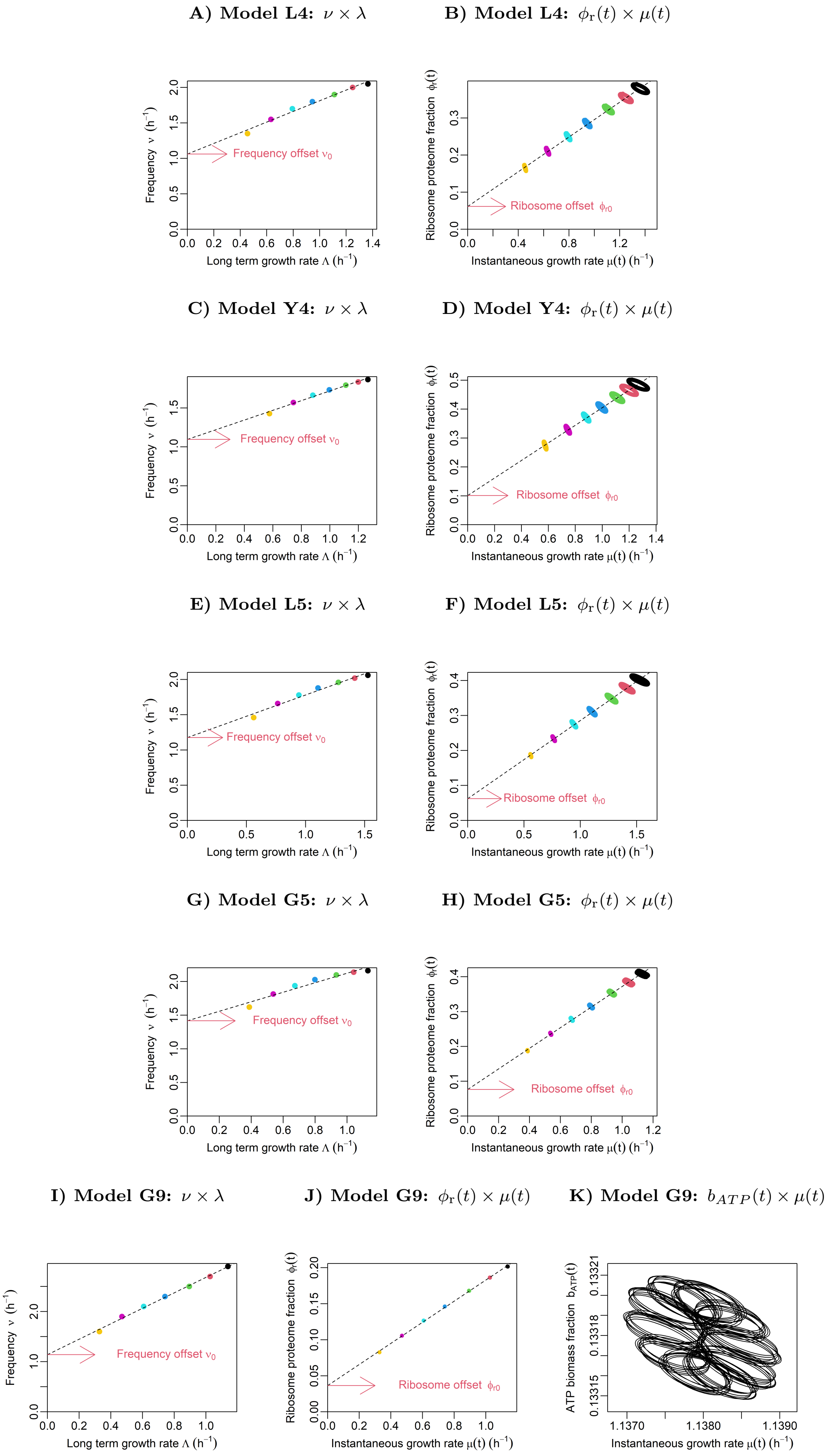


FIG. S20: Emergent almost linear relationships between optimal cell properties at different static media (different colors) as for model L3 on Fig.4 in the main text, here for models L4, Y4, L5, G5, G9.

#### Growth law for protein allocation

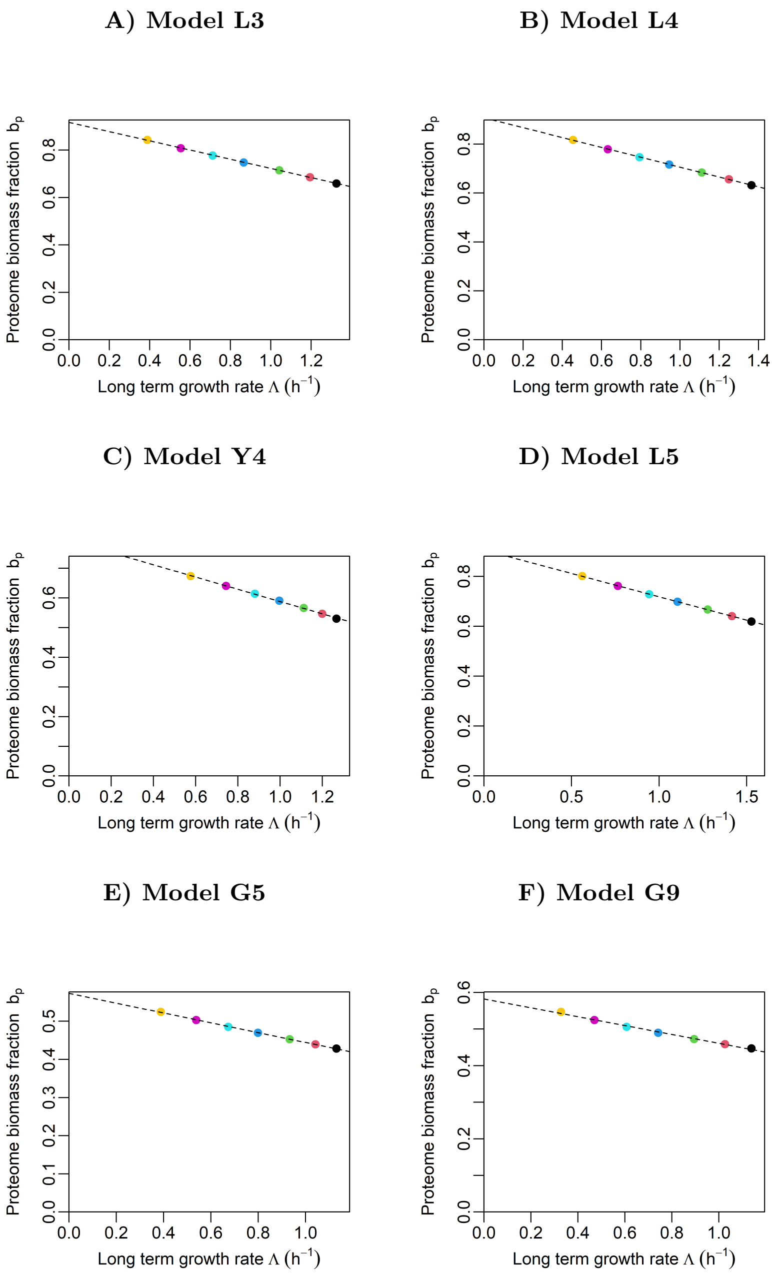


FIG. S21: The predicted proteome biomass fraction decreases almost linearly with the long term growth rate $\Lambda$ at different media, for all CGMs tested.

[1] W. Liebermeister and E. Klipp, Theoretical Biology and Medical Modelling **3**, 41 (2006), ISSN 1742-4682, URL https://doi.org/10.1186/1742-4682-3-41.
